# Lipid Droplet Is an Ancient and Inheritable Organelle in Bacteria

**DOI:** 10.1101/2020.05.18.103093

**Authors:** Xiang Chi, Ololade Omolara Ogunsade, Ziyun Zhou, Zemin Li, Xuehan Li, Mengwei Zhang, Fuhang Song, Jun Wang, Mirza Ahmed Hammad, Xuelin Zhang, Shuyan Zhang, Xia Wan, Lixin Zhang, Congyan Zhang, Pingsheng Liu

**Affiliations:** National Laboratory of Biomacromolecules, CAS Center for Excellence in Biomacromolecules, Institute of Biophysics, Chinese Academy of Sciences, Beijing 100101, China; University of Chinese Academy of Sciences, Beijing 100049, China; School of Kinesiology and Health, Capital University of Physical Education and Sports, Beijing 100191, China; CAS Key Laboratory of Pathogenic Microbiology and Immunology, Institute of Microbiology, Chinese Academy of Science, Beijing 100101, China; Oil Crops Research Institute of the Chinese Academy of Agricultural Sciences, Wuhan 430062, China; Key Laboratory of Biology and Genetic Improvement of Oil Crops, Ministry of Agriculture, Wuhan 430062, China; Oil Crops and Lipids Process Technology National & Local Joint Engineering Laboratory, Wuhan 430062, China; Hubei Key Laboratory of Lipid Chemistry and Nutrition, Wuhan 430062, China; State Key Laboratory of Bioreactor Engineering, and School of Biotechnology, East China University of Science and Technology (ECUST), Shanghai 200237, China; Department of Molecular and Cell Biology, University of California, Berkeley, Berkeley, California 94720, USA

**Keywords:** bacteria, lipid droplet, evolution, conservation, the last universal common ancestor with lipid droplet (LUCALD)

## Abstract

Lipid droplet (LD) is a monolayer phospholipid membrane-bound organelle found in all eukaryotes and several prokaryotes which plays key roles in cellular lipid homeostasis and human health. The origin and evolution of the organelle remains unknown. Here, we report that through screening over 660 bacteria using biophysical and biochemical methods, plus LD isolation and proteomic tool, LDs were identified in most of these microbes, affiliated with five main bacterial phyla. Moreover, LDs were also identified in *E. coli* overexpressing lipid synthesis enzymes, indicating that bacteria without detectable LDs possessed the ability of LD biogenesis. The similarity of isolated LDs from representative strains and evolutionary analysis of LD major protein PspA demonstrate that LDs were conserved in bacteria. Furthermore, time-lapse imaging revealed that LDs were inheritable accompanying with bacterial growth and division. Finally, a common ancestor of LD-containing bacteria was predicted to originate 3.19 billion years ago by a phylogenetic analysis. Our findings suggest that LD is a widespread and inheritable organelle from an ancient common ancestor.

## Introduction

Lipid droplet (LD) is a monolayer phospholipid membrane-bound organelle that has been found in almost all eukaryotic cells and plays multiple functions besides storage of neutral lipids [1-3]. LDs in eukaryotic cells are essential for the storage and production of food oil, nutrients, vitamins, biofuels, and other hydrophobic carbon sources [1]. In addition, many health conditions and disorders have been linked to as well as depended on or even caused by the homeostasis of this unique organelle in humans [4, 5]. The recent developments in new methods such as isolation and imaging, as well as omics studies including genomics, proteomics, lipidomics, and metabolomics have dramatically promoted the LD research [6] and in turn, generating much more new information. As a newly valued-organelle even discovered long ago, many fundamental knowledges about the organelle especially its evolution path are still missing and waiting to be discovered. Based on recent identified LD proteomes and related functional studies, roles of LDs have been proposed with some verifications, which includes but not limited to neutral lipid metabolism and storage [7, 8], protein storage and degradation [9, 10], intracellular transportation [11, 12], cytotoxicity reduction [13-15], and nucleic acid handling [16, 17]. Due to the similarity of some functions between LD and endoplasmic reticulum (ER), the terminal proteins in neutral lipid synthesis largely in the ER, as well as the physic contact between these two organelles, currently LD is proposed to be generated in the ER [1, 2]. But the facts that except cytoplasm LDs, the organelle has also been found in several bacteria [18-20] that do not have any ER or ER like structures, and more interestingly inside of other cellular organelles such as endoplasmic reticulum (ER), nucleus, chloroplast, and mitochondrion [21-27], suggesting another possibility that the LD is inheritable, and the origin of the organelle is from prokaryotic cells [28].

Eukaryotic LDs contain different neutral lipids such as triacylglycerol (TAG), cholesterol ester (CE)/ Sterol ester (SE), retinoid ester (RE), and ether lipid [29, 30]. Except for TAG, prokaryotes also have other neutral lipids such as wax ester (WE) and polyhydroxyalkanoates (PHA) [31]. Since these neutral lipids are hydrophobic molecules, they must gather together and are covered by a membrane structure, which allows them stay in aqueous phase cytoplasm. Exposure of neutral lipids to cytoplasm can be toxic to the cells. Therefore, as a result of a long evolutionary process the neutral lipids must have been covered by a certain amphipathic membrane, in order to distribute inside of cells and avoid cytotoxicity. Current finding on neutral lipids in eukaryotic cells is that all these neutral lipids are stored in LDs that are covered with a monolayer phospholipid membrane, a selective barrier, similar with other membranes, in which the hydrophobic acyl-chains of phospholipid face to hydrophobic neutral lipids and their hydrophilic head groups to aqueous phase of cytoplasm. It is possible that same with eukaryotes, prokaryotic cells may also store their neutral lipids in such structures.

Emerging of bilayer phospholipid membrane-bound organelles was proposed to meet the requirement of more bioenergetical efficiency. But this is challenged by recent lines of evidence that eukaryotic organisms are not with higher energetic capacity than prokaryotes [32]. In addition, prokaryotes have plasma membrane, a bilayer membrane, which has similar properties of bilayer membrane-bound organelles. If the plasma membrane is wrinkled toward to cytoplasm, it can generate more membrane area with curvature that provides solid surface for chemical reactions, which also challenges the necessity for higher reaction efficiency arose from bilayer membrane-bound organelles. The rational explanation is that prokaryotes evolved not only neutral lipids but also a monolayer membrane structure for storage of neutral lipids, which provides a higher density energy reservoir, a carbon source, and a catalytic surface for higher reaction efficiency.

Furthermore, this specialized monolayer membrane provides some unique properties. For example, some LD-associated proteins only localize on LDs and cannot be found in other bilayer membrane structures. These proteins are termed as LD resident proteins. In addition, this recognition for LD monolayer membrane can be achieved cross organisms [6, 33]. For example, exogenously expressed human LD resident protein, adipose differentiation-related protein (ADRP)/perilipin2 (PLIN2), is also localized on bacterial LDs, while worm LD resident mediator-28 (MDT-28) can localize on LDs in both mammalian cells and bacteria. Furthermore, three LD resident proteins including human ADRP/PLIN2, worm MDT-28, and bacterial microorganism lipid droplet small (MLDS) can be recruited to adiposomes, an artificial structure similar to LD with a TAG core covered by a monolayer phospholipid membrane [34], suggesting the conservation of this unique property of monolayer membrane.

To identify if bacteria have LDs, our group established a method to isolate those structures from bacteria and analyzed them morphologically and biochemically [35]. Our results identified LDs in a couple of bacteria and further found those LDs not only store neutral lipids as TAG with monolayer phospholipid membrane [19, 35] but also bind and protect genomic DNA on their surface through the LD resident protein MLDS [17]. After infection, hepatitis C virus (HCV) utilizes LD surface and ER of hepatocytes for its replication and assemble [36, 37], suggesting LD ability of nucleic acid handling. These findings indicate that the property of LD handling nucleic acids is conserved from bacteria to mammals [28].

LDs have been found in almost all eukaryotic cells with similar shape, contents, and properties, demonstrating that the organelle is conserved in Eukarya domain [38]. Several types of neutral lipids have been identified in prokaryotes including TAG [39]. In addition, the homologues of metabolic machineries of neutral lipids have also been found in many prokaryotic organisms [31, 40]. Therefore, this monolayer membrane-bound organelle may also be conserved and well distributed in prokaryotes. Searching for LDs in Bacteria domain may help us to determine the origin of this monolayer membrane-bound organelle and also enable us to explore the evolution of other membrane-bound organelles.

By screening more than 2,000 soil bacterial strains, we identified 660 individual strains and found that most of them had certain level of neutral lipids, and their distribution was similar with soil bacterial population. We further verified that these neutral lipids were stored in LDs through morphological studies, biochemical studies, as well as proteomics studies on LDs isolated from some representative strains. We also found that a common ancestor of LD-containing bacteria was evolved 3.19 billion years ago based on a phylogenetic analysis.

## Materials and Methods

### Bacterial strains

Bacteria used in this study include 2,000 soil strains and 10 commercial strains. The 2,000 soil strains (LS130001-LS132000) were obtained from the microbial culture library constructed by Lixin Zhang’s laboratory (Institute of Microbiology, Chinese Academy of Sciences). The 10 commercial strains were purchased as following: *Rothia dentocariosa* (ATCC® 14189™), *Bifidobacterium longum* (ATCC® 15707™), *Corynebacterium striatum* (ATCC® 9016™), *Acinetobacter calcoaceticus* (CGMCC 1.6186), *Rhodococcus rhodochrous* (CGMCC 1.2348), *Acinetobacter baumannii* (CICC 10980), *Rhodococcus equi* (CICC 22955), *Streptococcus mutans* (CICC 10387), *Propionibacterium acnes* (CICC 10312), and *Nostoc punctiforme* (FACHB 252). The engineered *E. coli* strains 2119 and 2053 were constructed in previous study [41]. 25% glutaraldehyde solution, 8% paraformaldehyde solution (EM grade), Embed 812 kit, uranyl acetate and lead citrate were all purchased from Electron Microscopy Sciences (Hatfield, USA). pJAM2-*gfp* was constructed in previous report [17].

### Bacterial culture conditions

The 2,000 soil strains (LS130001-LS132000) were cultivated at 28°C in ISP2 medium (Yeast extract 4 g/L, Malt extract 10 g/L, Glucose 4 g/L, pH 7.0-7.2). To isolate a pure strain from the liquid culture, cells were streaked on the LB agar plates, and the plates were inverted cultivated at 28°C. *Bifidobacterium longum* was cultured anaerobically using the protocol in the American Type Culture Collection (ATCC). Other bacteria were cultivated aerobically in mineral salt medium (MSM) in Erlenmeyer flasks at 30°C with 0.5 g/L NH_4_Cl as a nitrogen source and 10 g/L gluconate sodium as a carbon source according to previous method [17].

### Strain identification

The single clone isolated from the soil strains was used as the template to amplify the 16S ribosomal RNA gene using the PCR with *TransStart*® FastPfu DNA Polymerase and primers F (AGAGTTTGATCCTGGCTCAG) and R (TACGGCTACCTTGTTACGAC). The procedures for thermal cycling were as follows: denaturation of the target DNA at 95°C for 10 min, followed by 30 cycles at 94°C for 1 min, annealing at 65°C for 1 min, and extension at 72°C for 1 min. After the last PCR cycle, the reaction mixture was held at 72°C for 6 min then cooled to 4°C. The 16S rDNA sequences of these strains were sequenced and checked for species specificity in the National Center for Biotechnology Information (NCBI) database with BLAST software. Sequences with the best match to a database sequence were considered for identifying their taxonomy.

### Confocal microscopy

As described in the previous study [17], cultivated bacterial cells were stained with LipidTOX Red (H34476) at 1:500 (v/v) in the dark for 30 min and then loaded on cover glasses, pretreated with poly-L-lysine (PB0589) for 30 min. The cover glasses were mounted on glass slides using antifade mounting medium (P0126) and imaged with Olympus FV1200 confocal microscope.

For time-lapse live imaging, the bacterium *Rhodococcus jostii* RHA1 stained by LipidTOX Red was grown on a gel surface (2% agarose dissolved in MSM), and the gel was covered by a cover glass. The bacteria were then observed with Olympus FV1200 confocal microscope at 30°C for several hours. Images were collected images at 10-min interval, and the representative images as indicated time were shown.

For three-dimensional (3D) imaging and tomography, the samples were prepared according to previous report [17], and bacteria were observed by Olympus FV1200 confocal microscope with 3D model. Images were collected images at 10-nm interval, and around 40 images in total for one field were collected. Then these images were analyzed using Imaris 8.1.2 software.

### Enzymatic analysis

Cultivated bacteria (1 mL) were centrifuged at 10,000*g* for 1 min. The collected cells were washed twice with l mL PBS, and then dissolved in 200-400 µL 1% Triton X-100 by sonication. The whole cell lysates were then centrifuged at 10,000*g* for 5 min at 4°C. The supernatant was collected into a new Eppendorf tube. The TAG content of the supernatant was measured using the Triglycerides Kit (GPO-PAP Method) (BioSino Bio-Technology and Science Inc., China). Protein concentration was quantified using the Pierce™ BCA Protein Assay Kit (Thermo, USA).

### Thin layer chromatography

The lipids of cultivated bacterial cells were extracted twice with a mixture of chloroform/methanol/PBS (2:1:1, v/v/v). The lipids of purified LDs were extracted by chloroform/acetone (5:7, v/v). The organic phases were collected and dried under a stream of nitrogen gas. The resulting lipids were dissolved in 100 μL, or less, chloroform and then subjected to thin layer chromatography (TLC) analysis using a silica gel plate. The plate was developed in a solvent system of hexane/diethyl ether/acetic acid (80:20:1, v/v/v) to separate neutral lipids. Then lipids on the plates were visualized by iodine vapor in a box.

### Isolation of lipid droplets

A previously described method [42] was used to isolate lipid droplets from bacteria. Briefly, bacteria cells were collected and resuspended in Buffer A (25 mM tricine, 250 mM sucrose, pH 7.8). The resuspended cells were homogenized by passing through a French press cell six times at 1,000-1,500 bar at 4°C. The homogenate was then centrifuged at 6,000*g* for 10 min at 4°C. The supernatant was the whole cell lysate (WCL) fraction. The supernatant (8 mL) was transferred into a SW40 tube with 4 mL buffer B (20 mM HEPES, 100 mM KCl, 2 mM MgCl_2_, pH 7.4) on top, and then the sample was centrifuged at 182,000*g* for 1 h at 4°C (Beckman SW40). LD fraction was carefully collected from the top band of the gradient and washed three times using 200 μL of buffer B. To prepare the LD protein sample, 1.2 mL of chloroform-acetone (5:7, v/v) was added to isolated LDs to precipitate proteins and extract lipids. LD proteins were dissolved in 20-50 μL 2×SDS sample buffer, and denatured at 95°C for 5 min.

### Mass spectrometry (MS) analysis

LD proteins were separated on a 10% or 12% SDS-PAGE gel and subjected to silver staining or colloidal blue staining. For in-gel digestion, each slice of the gel was cut and destained with 30 mM K_3_Fe(CN)_6_ and 100 mM Na_2_S_2_O_3_, and then dehydrated with 100% acetonitrile. Proteins were reduced with 10 mM DTT in 25 mM NH_4_HCO_3_ at 56°C for 1 h and alkylated by 55 mM iodoacetamide in 25 mM NH_4_HCO_3_ in the dark at room temperature for 45 min. Finally, gel pieces were thoroughly washed with 25 mM NH_4_HCO_3_, 50% acetonitrile and 100% acetonitrile respectively and completely dried in a SpeedVac. Trypsin was added to a ratio of 1:40 relative to total substrate and incubated overnight at 37°C. The digestion reaction was stopped by addition of formic acid to a final concentration of 0.5%. The gel pieces were extracted twice with 80 μL 60% acetonitrile plus 0.1% formic acid, and then sonicated for 15 min. All liquid samples from the two extractions were combined and dried in a SpeedVac. For in-solution digestion, the LD proteins were dissolved in suitable amount of 8 M urea, reduced with 10 mM DTT at room temperature for 1 h, and alkylated by 40 mM iodoacetamide in 25 mM NH_4_HCO_3_ in the dark at room temperature for 45 min. A second addition of DTT to a final concentration of 40 mM was made to neutralize any remaining iodoacetamide. The sample was then diluted with 25 mM NH_4_HCO_3_ to reduce the concentration of urea less than 2 M. Trypsin was added to a ratio of 1:50 relative to total substrate and incubated overnight at 37°C. The digestion reaction was stopped by addition of formic acid to give a final concentration of 0.5%. The sample was then centrifuged at 13,000*g* for 20 min and the supernatant was collected and stored at −20°C for further study.

Dried peptide samples were dissolved in 20 μL 0.1% formic acid, loaded onto a C18 trap column with an autosampler, eluted onto a C18 column (150 mm × 75 μm) packed with 3μm ReproSil-Pur C18-AQ (Dr. Maisch GmbH, Ammerbuch) packing material, and then subjected to nano-LC-LTQ-Orbitrap XL (Thermo, San Jose, CA) MS/MS analysis.

MS/MS data were searched using SEQUEST program (Thermo, USA) against the NCBI Refseq bacteria database. Search parameters were set as follows: enzyme cleavage specificity: trypsin; no more than two missed cleavages; precursor ion mass tolerance: 20 ppm; and fragment ion mass tolerance: 0.6 Da. The fixed modification was set to carboxyamidomethylation of cysteine. The variable modification was set to oxidation of methionine. The SEQUEST outputs were then analyzed using the software Proteome Discoverer (version 1.4.0.288, Thermo Fischer Scientific). The peptide-spectrum matches (PSM) were filtered using the Percolator algorithm, and the q value is less than 1% (1% FDR). The retrieved peptides were combined into proteins using strict maximum parsimony. At the protein level, only proteins with at least two unique peptides were accepted.

### Construction of green fluorescent fusion proteins

PspA was amplified without the native start and stop codons using the primers F (TTAGGATCCGCTAATCCTTTCGTCAAGGG) and R (TTAGGATCCCTGCCCGGTCTGACCGGCAG). The truncation was cloned into the BamHI site of pJAM2-*egfp*.

### Transmission electron microscopy

The ultra-structure of bacterial cells was examined using transmission electron microscopy (TEM) after ultra-thin sectioning. In brief, bacterial cells were collected and washed twice with 50 mM K-Pi (pH 7.2). Then the cells were fixed in 50 mM K-Pi containing 2.5% (v/v) glutaraldehyde and 2% (v/v) paraformaldehyde overnight at 4°C. The cells were subsequently fixed in 2% (w/v) potassium permanganate for 5 min at room temperature. After dehydrated in an ascending concentration series of ethanol at room temperature, the samples were embedded in Embed 812 and prepared as 70 nm sections using Leica EM UC6 Ultramicrotome. Ultrathin sections were mounted on formvar copper grid and stained with uranyl acetate and lead citrate. The sections were then observed with Tecnai Spirit electron microscope (FEI, Netherlands).

The isolated lipid droplets were also examined by TEM through negative staining. Briefly, the isolated lipid droplets were placed on a formvar copper grid and stained for 30 s by 2% (w/v) uranyl acetate. The grid was then viewed with Tecnai Spirit electron microscope (FEI, Netherlands).

### Phylogenetic analysis

The bacteria containing LDs in this study and previous reports were analyzed by Timetree database [43]. Then the evolutionary tree was displayed via Interactive Tree Of Life [44]. Different colors represented different phyla. The estimated time of each node in the evolutionary tree was indicated.

## Results

### Many bacteria contain neutral lipids including triacylglycerol

Almost all eukaryotic cells have been found to contain or be able to form LDs [28, 38]. Although neutral lipid synthetic enzymes have been found in many bacterial strains [45], LDs or LD-like structures (lipid inclusions) have been visualized and systematically analyzed in only few bacterial strains (Fig. S1A) [38]. To explore whether this organelle exists in most bacteria, more than 2,000 bacterial isolates were collected from soils in several locations in China and neutral lipids and LDs were examined (Fig. S1B). Among them, 660 individual stains were determined through 16S rRNA gene amplicon sequencing (Table S1). These bacteria mainly consisted of four phyla, including *Actinobacteria* (45.0%), *Proteobacteria* (33.5%), *Firmicutes* (17.0%), and *Bacteroidetes* (2.3%) (Fig. 1A, left panel). The right panel of Figure 1A presented major genera of these abundant phyla. Moreover, *Actinobacteria* and *Proteobacteria* were the major bacterial phyla in this collection, similar to the composition of dominant soil bacteria reported before [46], suggesting that the collection is representative for soil bacteria. Since neutral lipids are stored in LDs and TAG is a major neutral lipid in LDs of eukaryotic cells, we first screened the 660 strains for TAG content using enzymatic analysis. The commercial *Escherichia coli* (*E. coli*) strain TOP10 was used as a negative control since the LD is not detected in the bacterium. The TAG content of *E. coli* strain TOP10 was measured as 3 (μg/mg protein) (Table S2). The *Mycobacterium smegmatis* and *Rhodococcus jostii* RHA1 that are previously reported to have TAG-containing LDs were used as positive controls [35, 47] and their TAG contents were about 100 and 700 (μg/mg protein), respectively (Table S2 and Fig. 1Ba). Figure 1Ba presented some of these results. Therefore, 100 (μg/mg protein) was used as middle level and 700 (μg/mg protein) used as high level (Fig. 1Ba). We found that the ratio of bacteria with TAG content equal to or less than 3 (μg/mg protein) was about 1.4%, indicating that these bacteria may not contain LDs (Figure 1Bb, Table S2). The ratio of bacteria with TAG content more than 3 and less than 100 (μg/mg protein) was about 41.1%, indicating that these bacteria have the possibility of containing LDs (Figure 1Bb, Table S2). The ratio of bacteria with TAG content between 100 to 700 (μg/mg protein) or more than 700 (μg/mg protein) were about 54.4% and 3.2%, respectively, suggesting that these bacteria (at least 57.6%) should contain LDs (Figure 1Bb, Table S2).

**Figure 1.**
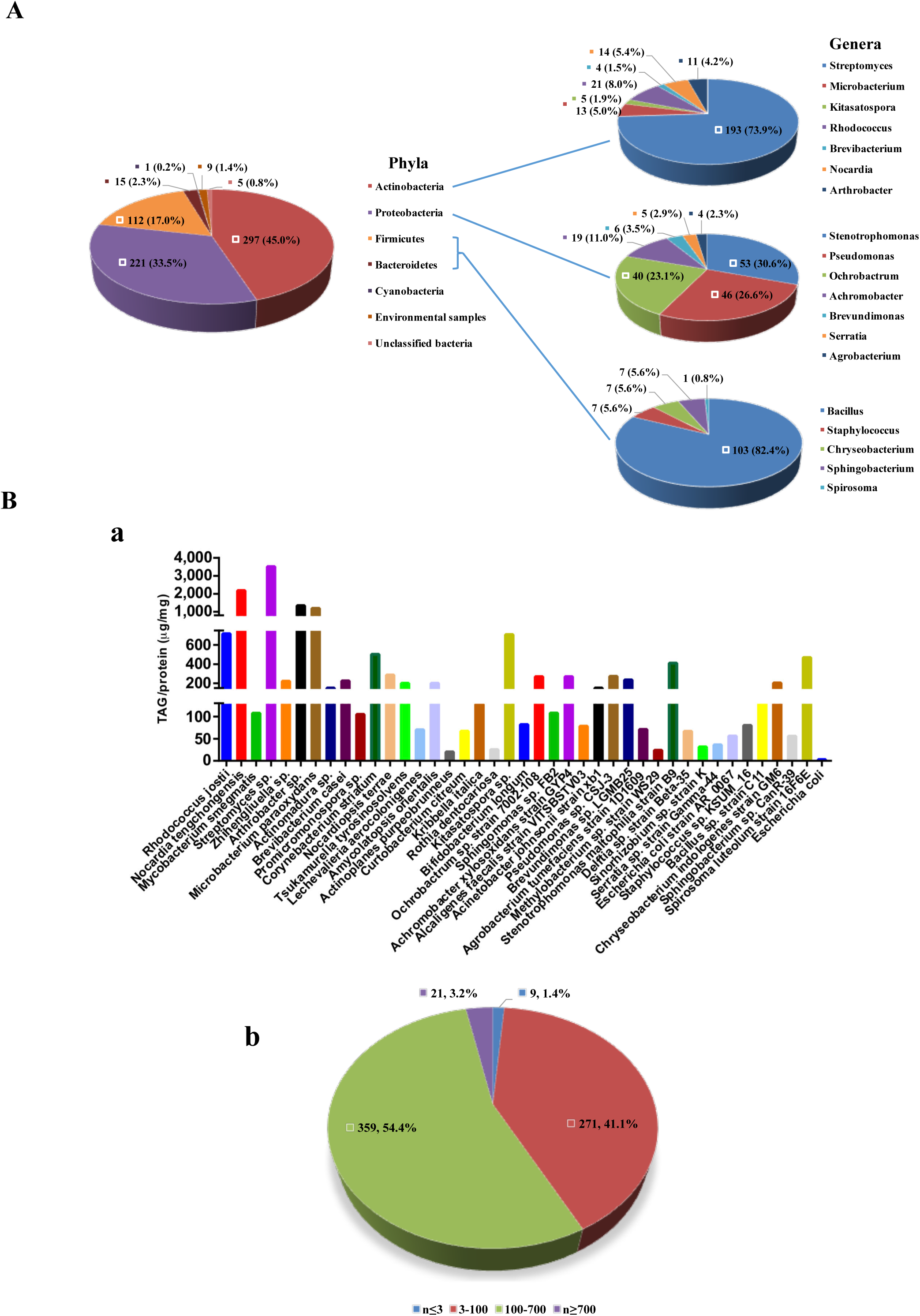
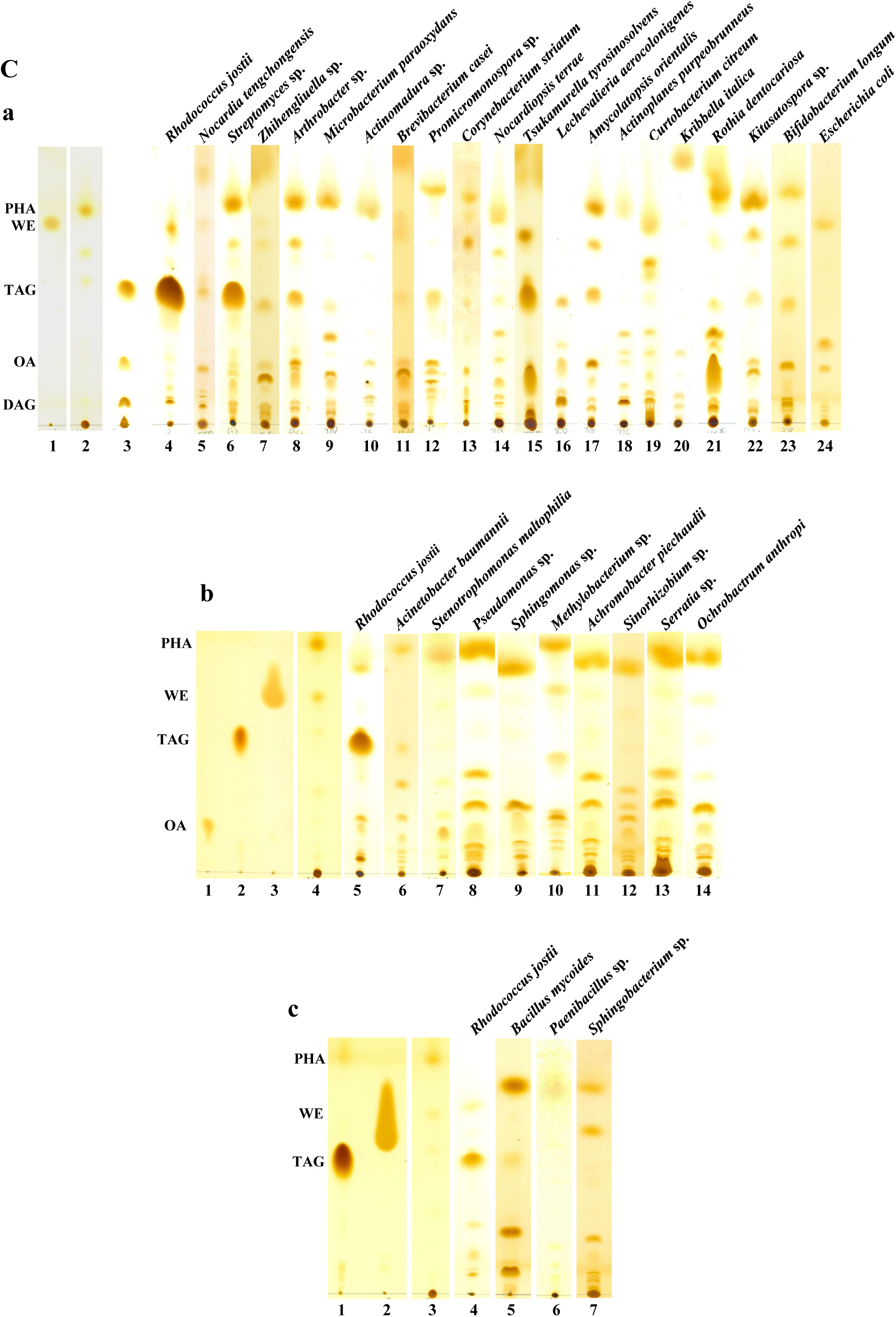
Neutral Lipid Screening of Bacteria. Bacterial strains were cultivated in the conditions described in Materials and Methods. Total lipids were extracted using Bligh-Dyer method, triacylglycerol (TAG) was measured by enzymatic assay, and neutral lipids were separated and determined by thin layer chromatography (TLC). Lanes 1-3 are the standards. From top to bottom: polyhydroxyalkanoate (PHA), wax ester (WE), triacylglycerol (TAG), oleic acid (OA), and diacylglycerol (DAG). **A** The distribution of the identified bacteria in different phyla (left panel) and major genera of these abundant phyla (right panel). **B a** A representative figure that shows TAG contents in selected bacterial strains. **b** The distribution of the strains with different TAG levels. **C** Representative TLC analyses of the selected strains. **a** Selected strains of *Actinobacteria*. **b** Selected strains of *Proteobacteria*. **c** Selected strains of *Firmicutes* and *Bacteroidetes*.

To determine other neutral lipids such as WE and PHA, based on the abundance of each genus, representative strains identified by the 16S rRNA sequencing method in each genus were randomly selected according to their genera abundance to provide sufficient detection range of diverse neutral lipids contained in each genus. Total lipids were extracted and subjected to thin layer chromatography (TLC) for separation and detection. Figure 1C presented representative data for some strains selected from phyla *Actinobacteria* (Fig. 1Ca), *Proteobacteria* (Fig. 1Cb), and *Firmicutes* and *Bacteroidetes* (Fig. 1Cc). The rest of TLC analyses for other strains was showed in Figure S2 and grouped by *Actinobacteria* (Fig. S2Aa), *Proteobacteria* (Fig. S2B), *Firmicutes* (Fig. S2C), and B*acteroidetes* (Fig. S2D). Based on these TLC analyses, for 58 strains of *Actinobacteria*, 67.2% contained TAG, 72.4% contained WE, and 100% contained PHA (Table S3). These numbers were 11.2% (TAG), 60.2% (WE), and 96.9% (PHA) among 98 strains of *Proteobacteria*. There were 3 TAG positive, 17 WE positive, and 19 PHA positive in 21 strains of *Firmicutes*, and 1 for TAG, 1 for WE, and 2 for PHA in 4 strains of *Bacteroidetes* (Table S3). Importantly, all these analyzed strains from four major phyla of bacteria were found to have certain types of neutral lipids, suggesting the possibility that these bacteria have LDs or similar structures for neutral lipid storage.

### Neutral lipid-containing bacteria have lipid droplet-like structures

Based on the principle “like dissolves like”, the hydrophobic neutral lipids should be sequestered together and covered by an amphipathic membrane, phospholipid monolayer, in hydrophilic cytoplasm. Those structures in cells are, in fact, an organelle, lipid droplet. To determine whether the bacterial strains positive for neutral lipids contain LDs, some of representative strains were selected, stained by LipidTOX and imaged using confocal microscopy. Some of them were analyzed using transmission electron microscopy (TEM) for further verification. Bacterial strains of phylum *Actinobacteria* were examined first because they are major bacteria in our collection and it has been reported that a couple of *Actinobacteria* strains contain LDs [38]. This study further shows that many strains of *Actinobacteria* contain LipidTOX Red-positive and LD-like structures (Fig. 2A and Fig. S3A). Next, the bacteria from other major phyla in our collection were visualized using the same methods, including *Proteobacteria, Firmicutes*, and *Bacteroidetes*. The strains containing LipidTOX Red-positive and LD-like structures were presented in Figures 2B and S3B (*Proteobacteria*), Figures 2C and S3C (*Firmicutes*), and Figures 2D and S3D (*Bacteroidetes*). Together, 79 strains of *Actinobacteria*, 19 strains of *Proteobacteria*, 8 strains of *Firmicutes*, and 3 strains of *Bacteroidetes* were found to have LipidTOX Red-positive and LD-like structures (Table S4). One strain of *Cyanobacteria, Nostoc punctiforme*, was stained with BODIPY (Fig. S3E, Table S4) [48]. To verify these LipidTOX Red-positive and LD-like structures in the bacteria, some of these strains were selected and subjected to TEM analyses. LD-like structures could be detected in these representative strains from all four phyla by TEM (Fig. 2E). *Rhodococcus jostii* RHA1 from *Actinobacteria* represented the bacterial strain in which TAG was a major neutral lipid (Fig. 1C, lane 4, Fig. 2Ec and 2Ed), while *Brevibacterium casei* strain KA2 (2Ee and 2Ef) and *Amycolatopsis orientalis* strain JAR10 (Fig. 2Eg and 2Eh), also from phylum *Actinobacteria*, represented the strains that mainly contained TAG and other neutral lipids (Fig. 1Ca, lane 11 and Fig. S2Ac, lane LS131278). The rest of strains such as *Sinorhizobium* sp. K (Fig. 2Ei and 2Ej) and *Ochrobactrum anthropi* TJ-1-58 (Fig. 2Ek and 2El) from phylum *Proteobacteria, Bacillus mycoides* WAB2225 (Fig. 2Em and 2En) from *Firmicutes*, and *Sphingobacterium ginsenosidimutans* (Fig. 2Eo and 2Ep) from *Bacteroidetes* were the bacteria that majorly contained WE and PHA. Figure 2 shows that neutral lipid-positive bacteria had LipidTOX Red-positive and LD-like structures, without neutral lipid specificity, suggesting that neutral lipids in bacteria, similar to eukaryotes, are stored in LD-like structures following the principle “like dissolves like”.

**Figure 2.**
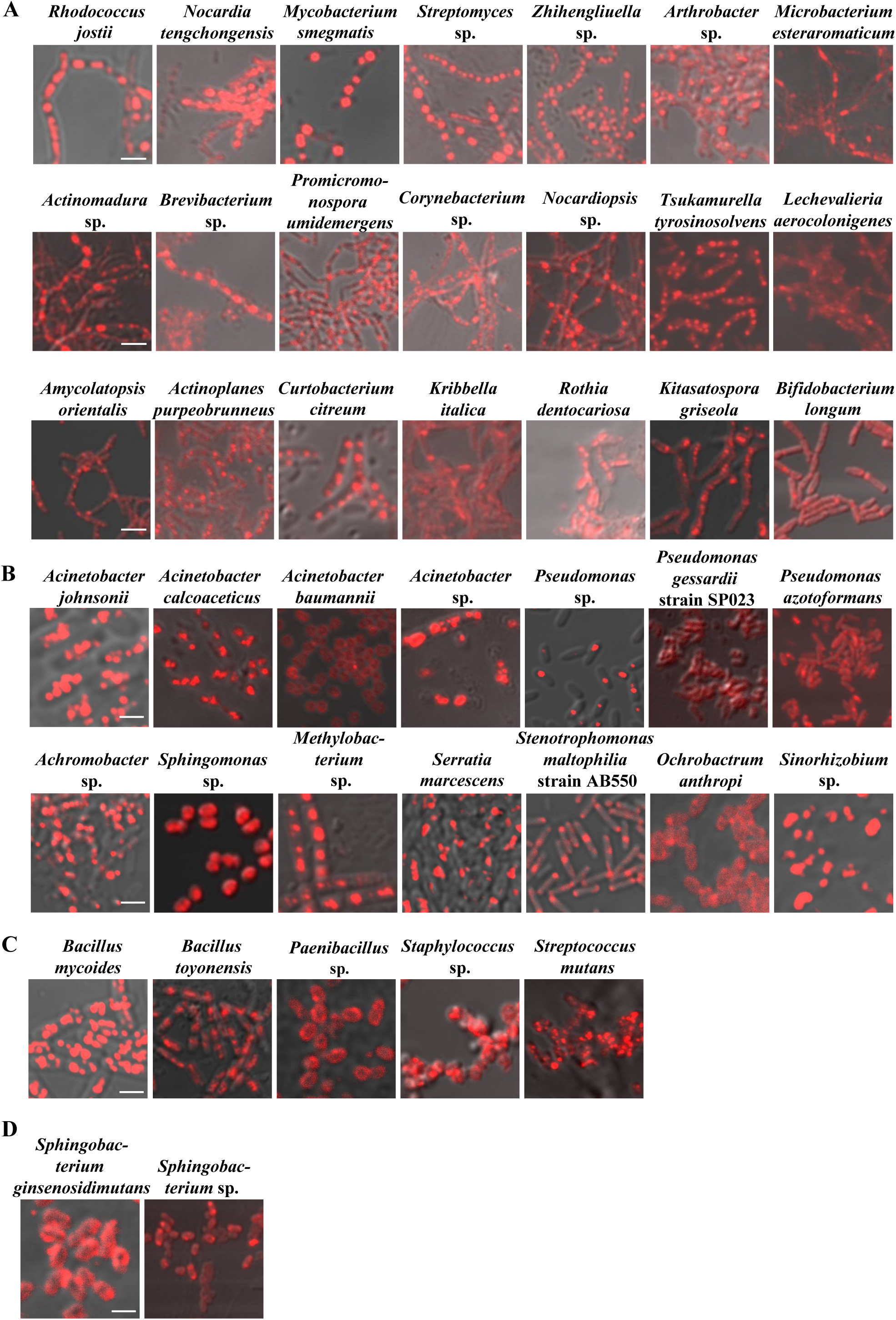
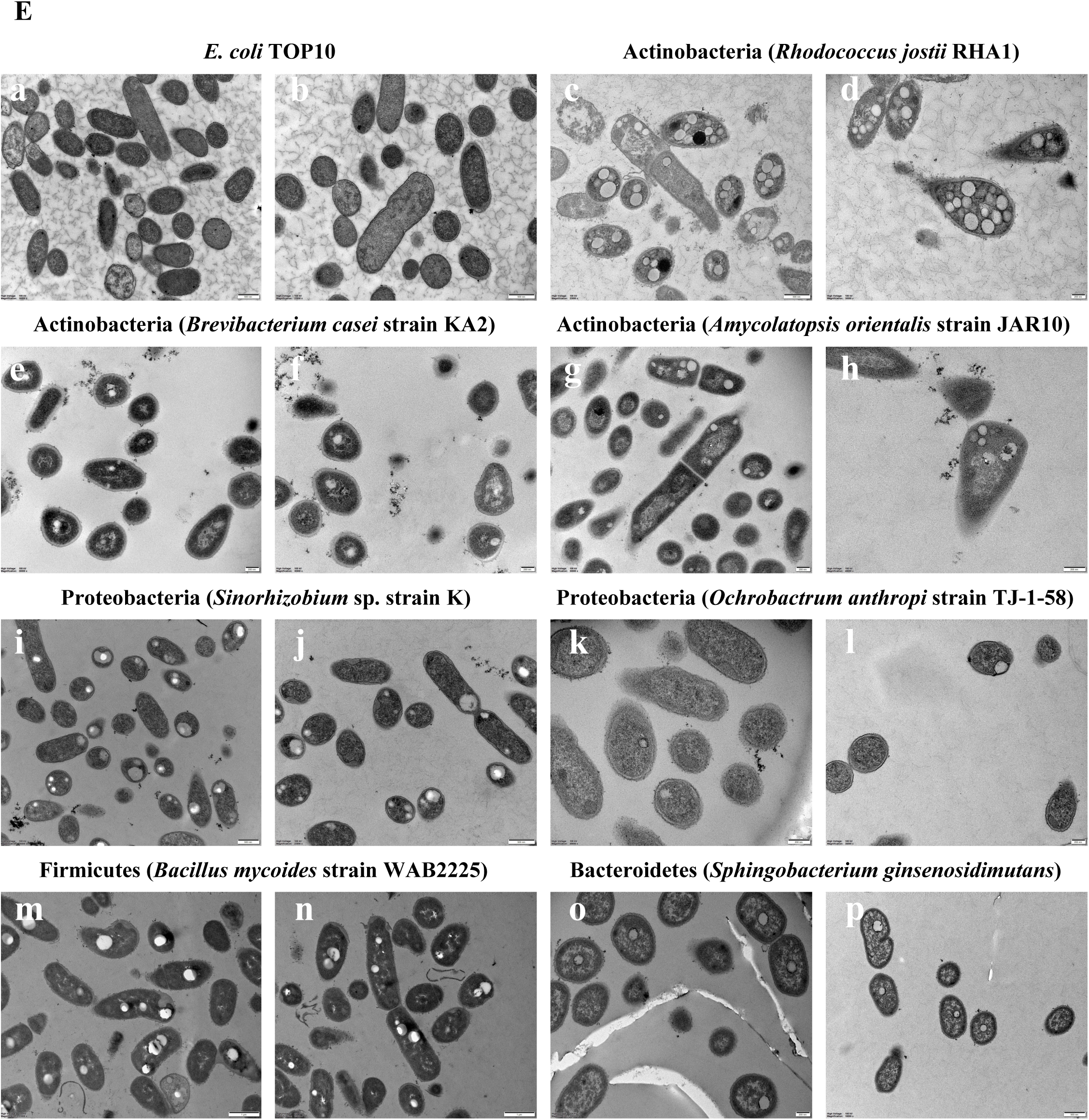
Morphological Screening of Neutral Lipid-containing Bacteria. Bacterial strains were cultivated in the conditions described in Materials and Methods. Neutral lipids in these strains were stained by LipidTOX Red and imaged using confocal microscopy. Scale bar = 2 µm. Lipid droplet (LD)-like structures were determined using transmission electron microscopy (TEM). **A** The staining of neutral lipids in representative strains of *Actinobacteria*. **B** The staining of neutral lipids in representative strains of *Proteobacteria*. **C** The staining of neutral lipids in representative strains of *Firmicutes*. **D** The staining of neutral lipids in representative strains of *Bacteroidetes*. **E** The ultra-structure of the representative strains was observed using TEM. Briefly, the cells were dehydrated through an ethanol series after a two-step fixation. The samples were then embedded in resin and thin sectioned. The sections were stained and visualized by the TEM. In **a**-**c, i, j**, and **p**, Scale bar = 500 nm. In **d**-**h, k, l** and **o**, Scale bar = 200 nm. In **m** and **n**, Scale bar = 1 μm.

### Lipid droplet-like structures are isolated and confirmed to be lipid droplets

To demonstrate these LD-like structures in bacteria are indeed the organelle, LDs, these structures have to be isolated and analyzed to determine if they fit the LD criterions. Based on current understandings, LD is defined as a cellular organelle with the features: 1) spherical shape, 2) phospholipid monolayer membrane and neutral lipid core, 3) associated proteins especially LD resident proteins, and 4) functions. We established the methods to isolate LDs from bacteria and analyzed them using morphological, biophysical, and biochemical means near a decade ago [35], which allows us to systematically study bacterial LDs in a higher quantity. Thus, LDs were isolated from representative bacterial strains of major phyla of our collection. The isolated LDs were then analyzed at four levels: 1) LipidTOX Red-staining structure to overlap with and imaging using confocal microscopy, 2) size measurement using Delsa Nano C particle analyzer, 3) neutral lipid detection using TLC, and 4) proteomic study using LC/MS/MS. Figure 3A shows isolated LDs from six strains of *Actinobacteria*. The spherical shaped structures were stained well with LipidTOX Red, suggesting that they were relatively pure LDs (Fig. 3A). The size was then measured for other three isolated LDs from phyla *Actinobacteria*. Their average sizes were 195.1 nm (*Amycolatopsis*), 183.4 nm (*Nocardia*), and 645.2 nm (*Rhodococcus*), respectively (Fig. 3Bb, Fig. 3Cb, and Fig. 3Db). TLC data of neutral lipids from these three bacterial LD fractions shows that *Amycolatopsis* and *Nocardia* had TAG and other neutral lipids, and only TAG was detected in *Rhodococcus*. The initial spots of TLC plate suggested a low ratio of phospholipids (Fig. 3Bc, lane 2; Fig. 3Cc, lane 2; and Fig. 3Dc, lane 2) [29]. Total LD-associated proteins from *Rhodococcus opacus* identified by MS were classified into 9 categories and mainly included proteins of metabolism-related, ribosome, transcription- and translation-related, cell division-related, stress response, and chaperone (Fig. 3Dd). Compared with whole cell lysate (WCL), cytosol (Cyto), and total membrane (TM), isolated LDs (LD) had a unique protein composition (Fig. 3De, lanes 1-4) and the major protein bands in LDs were sliced and subjected to MS analysis, and identified proteins were listed on the right side (Fig. 3De). In agreement with previous study [35] MLDS and phage shock protein A (PspA) were identified from two major bands, RO3 and RO4 (Fig. 3De).

**Figure 3.**
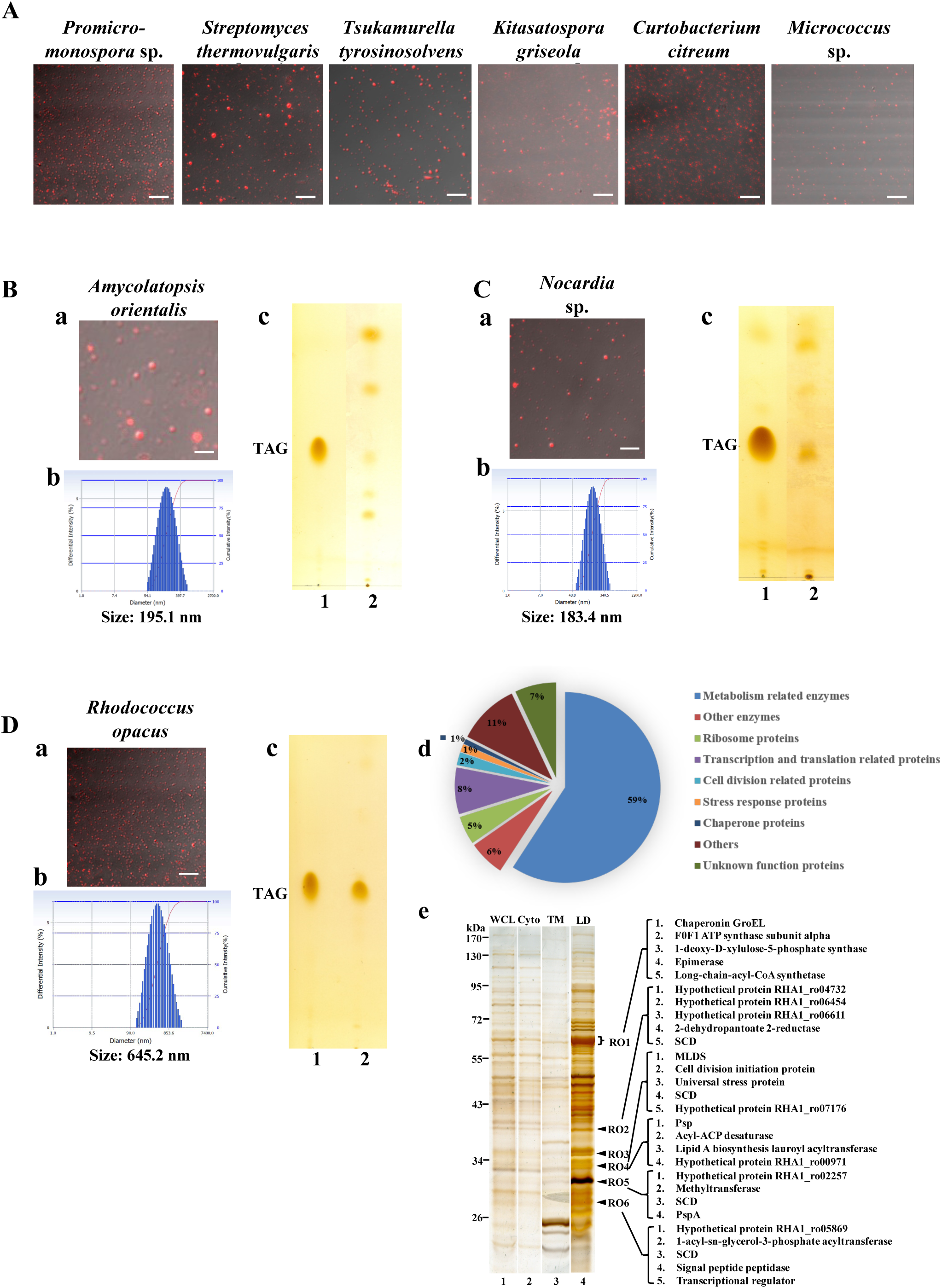
Isolation and Analysis of Lipid Droplets from Phylum *Actinobacteria*. Bacterial strains were cultivated in the conditions and lipid droplets (LDs) were isolated as described in Materials and Methods. The isolated LDs were stained by LipidTOX Red and imaged using confocal microscopy. Their lipids were extracted, separated and determined using TLC. Their proteins were identified by proteomic studies. **A** LDs were isolated from representative strains of phylum *Actinobacteria*, stained by LipidTOX Red and imaged using confocal microscopy. Scale bar = 2 µm. *Promicro., Promicromonospora*. **B** and **C** LDs were isolated from *Amycolatopsis* and *Nocardia*. **a** The isolated LDs were stained by LipidTOX Red and imaged using confocal microscopy. Scale bar = 2 µm. **b** The size of isolated LDs was measured using Delsa Nano C particle analyzer. **c** The neutral lipids of isolated LDs were analyzed using TLC. **D** LDs were isolated from a bacterium *Rhodococcus*. **a** The isolated LDs were stained by LipidTOX Red and imaged using confocal microscopy. Scale bar = 2 µm. **b** The size of isolated LDs was measured using Delsa Nano C particle analyzer. **c** The neutral lipids of isolated LDs were analyzed using TLC. **d** The whole proteins from isolated LDs were identified using shotgun mass spectrometry analysis and the identified proteins were categorized by their functions. **e** The whole proteins from isolated LDs (LD) were separated by SDS-PAGE, silver stained, and compared with whole cell lysate (WCL), cytosol (Cyto), and total membrane (TM). Then the major protein bands in LDs were sliced and subjected to mass spectrometry analysis, and identified proteins were marked.

To extend our knowledge and understanding of bacterial LDs, LDs from more bacterial strains of phylum *Actinobacteria* were studied by LD isolation and further analyses. First, LDs were also isolated from one strain of *Actinobacteria, Mycobacterium smegmatis*. The isolated LDs were also analyzed of morphology (Fig. S5Aa and b), size (Fig. S5Ac), neutral lipids (Fig. S5Ad), and protein composition (Fig. S5Ae and f). LDs from other strains of *Actinobacteria* were also isolated and analyzed of morphology, size, and neutral lipids, including gut bacteria *Bifidobacterium* (Fig. S4Aa-c), *Streptomyces* (Fig. S4Ba-c), *Kitasatospora* (Fig. S4Ca-c), and *Tsukamurella* (Fig. S4Da-c). LDs from *Amycolatopsis orientalis* of *Actinobacteria* were isolated and analyzed of morphology, size, and protein composition that was not only compared with other cellular fractions but also with LDs from *Rhodococcus jostii* RHA1 (Fig. S4Ga-c). LDs from more strains of *Actinobacteria* were also isolated and analyzed of morphology and protein composition, such as *Rhodococcus* sp. (Fig. S4Ha and b), *Streptomyces tanashiensis* (Fig. S4Ia and b), *Streptomyces mutomycini* (Fig. S4Ja and b), *Curtobacterium citreum* (Fig. S4Ka and b), and *Streptomyces griseus* (Fig. S4La and b). Three more LD isolations were carried out for phylum *Actinobacteria* and only analyzed of protein composition, including *Brevibacterium casei* and *Microbacterium esteraromaticum* (Fig. S4N), *Arthrobacter rhombi* (Fig. S4P), and *Actinomycete* HVG71 (Fig. S4Q). Compared with WCL, Cyto, and TM, the unique protein composition of the organelle is commonly used to determine the quality of its isolation [42].

Since neutral lipids and similar morphologies were also found in many strains of phylum *Proteobacteria*, the similar experiments were carried out on bacteria from this phylum. Isolated LDs from 4 strains were stained by LipidTOX Red and imaged using confocal microscopy, and the overlap verified LDs in *Proteobacteria* (Fig. 4A). Specifically, LDs were isolated from *Stenotrophomonas* sp. and stained with LipidTOX Red (Fig. 4Ba). Size of the LDs was measured (Fig. 4Bb) and their neutral lipids were analyzed using TLC (Fig. 4Bc, lane 2). Same as lipid composition of whole cell (Fig. 1Db, lane 7), the major neutral lipid in LDs was PHA (Fig. 4Bc, lane 2). The whole proteins of the isolated LDs were subjected to proteomic analysis and the identified LD-associated proteins were grouped into 6 categories. Three categories were similar to other LD proteomes, such as metabolism-related proteins, ribosome proteins, and DNA-binding proteins (Fig. 4Bd). In addition, protein composition of isolated LDs was distinct from other cellular fractions, such as WCL, Cyto, and TM. The main protein bands of LDs were subjected to MS identification and the determined proteins were listed on the right side of Figure 4Be. Several LD-associated proteins were found, including long chain fatty acid-CoA ligase (Fig. 4Be, band SM2), chaperone (Fig. 4Be, band SM2), and stomatin (Fig. 4Be, band SM5). The most abundant protein was PHA granule-associated protein (Fig. 4Be, band SM6).

**Figure 4.**
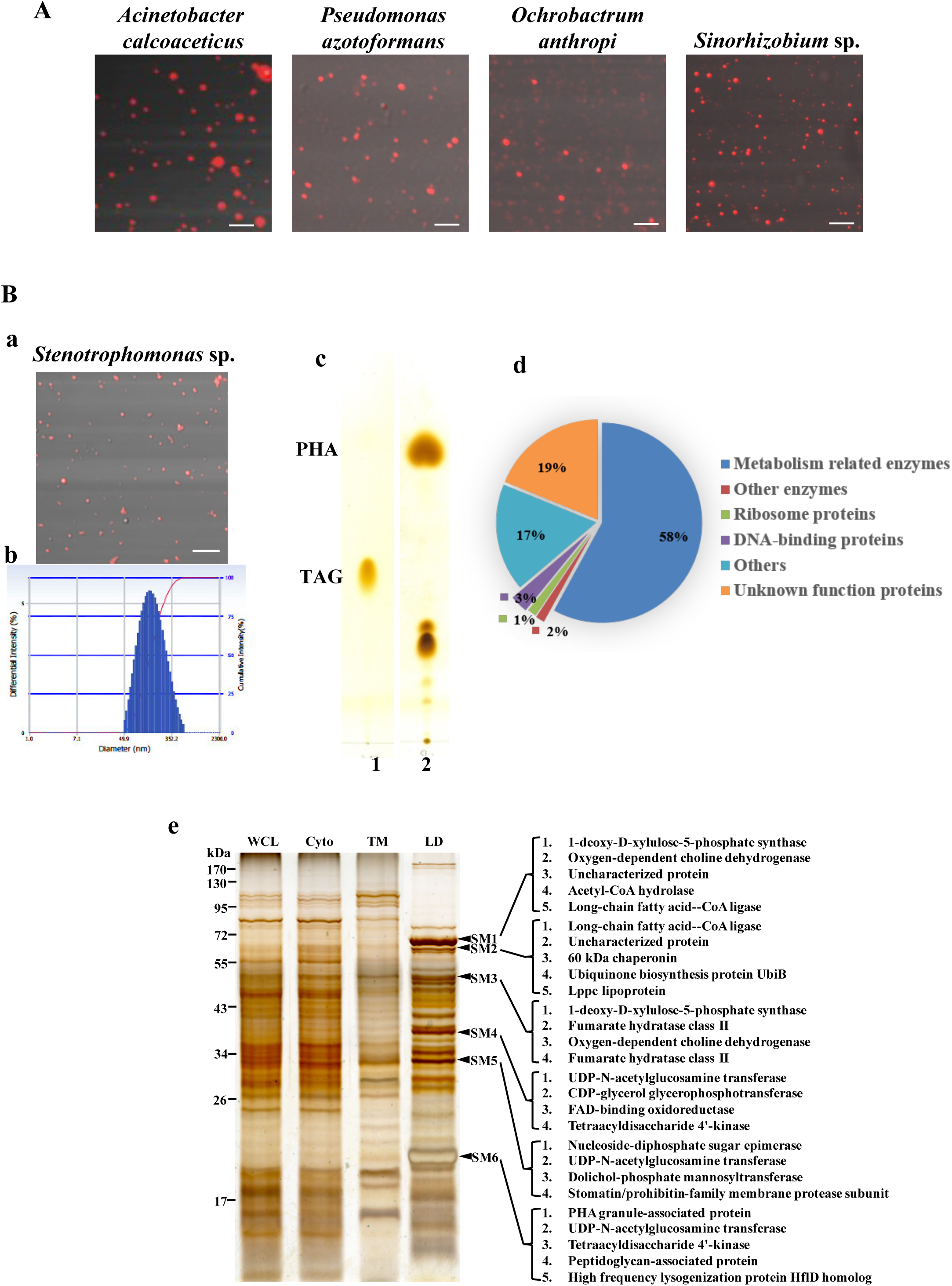
Isolation and Analysis of Lipid Droplets from Phylum *Proteobacteria*. Bacterial strains were cultivated in the conditions and lipid droplets (LDs) were isolated as described in Materials and Methods. The isolated LDs were stained by LipidTOX Red and imaged using confocal microscopy. Their lipids were extracted, separated and determined using TLC. Their proteins were identified by proteomic studies. **A** LDs were isolated from representative strains of phylum *Proteobacteria*, stained by LipidTOX Red and imaged using confocal microscopy. Scale bar = 2 µm. **B** LDs were isolated from a bacterium *Stenotrophomonas*. **a** The isolated LDs were stained by LipidTOX Red and imaged using confocal microscopy. Scale bar = 2 µm. **b** The size of isolated LDs was measured using Delsa Nano C particle analyzer. **c** The neutral lipids of isolated LDs were analyzed using TLC. **d** The whole proteins from isolated LDs were identified using shotgun mass spectrometry analysis and the identified proteins were categorized by their functions. **e** The whole proteins from isolated LDs (LD) were separated by SDS-PAGE, stained by silver staining, and compared with whole cell lysate (WCL), cytosol (Cyto), and total membrane (TM). Then the major protein bands in LDs were sliced and subjected to mass spectrometry analysis, and identified proteins were marked.

More LDs were isolated and analyzed from phylum *Proteobacteria*. In agreement with LipidTOX Red staining, BODIPY also labeled neutral lipids in bacterium *Sinorhizobium* sp. (Fig. S5Ba). Their LDs were then isolated. The isolated LDs were stained with LipidTOX Red (Fig. S5Bb), and their size was measured (Fig. S5Bc). The lipid composition was determined by TLC (Fig. S5Bd). The whole LD proteins were subjected to proteomic study and the identified LD proteins were classified into 9 categories and mainly included proteins of metabolism-related, ribosome, transcription- and translation-related, cell division-related, stress response, and chaperone (Fig. S5Be). The LD proteins were also separated by SDS-PAGE and analyzed by silver staining (Fig. S5Bf). The main LD protein bands were analyzed using proteomic tool and the proteins were listed in Figure S5Cf. LDs in another strain *Stenotrophomonas maltophilia* of phylum *Proteobacteria* were isolated and studied (Fig. S5C). First, the neutral lipid-related structure in the bacterium was stained using LipidTOX Red (Fig. S5Ca). Similar to other LD studies, briefly, LDs were isolated from *Stenotrophomonas maltophilia*, stained (Fig. S5Cb), size measured (Fig. S5Dc), neutral lipid analyzed (Fig. S5Dd), and protein determined (Fig. S5Ce and f). So far, LDs from three strains of phylum *Proteobacteria* were isolated and analyzed, respectively. Compared with phylum *Actinobacteria*, their main neutral lipids were not TAG. Therefore, these LDs were representative of the organelle containing other neutral lipids such as WE and PHA. In addition, LDs from four more strains of phylum *Proteobacteria* were also isolated, and two of them were analyzed of morphology and size, including *Pseudomonas* (Fig. S4Ea and b), and *Ochrobactrum* (Fig. S4Fa and b), while other two were only analyzed of protein composition, including *Acinetobacter calcoaceticus* (Fig. S4Ma and b) and *Acinetobacter baumannii* (Fig. S4O).

After extensively analyzing LDs in phyla *Actinobacteria* and *Proteobacteria*, we moved on to other two phyla *Firmicutes* and *Bacteroidetes*. Figure 5A presented results of isolated LDs from strain *Bacillus mycoides* of phylum *Firmicutes*. The isolated LDs were stained using LipidTOX Red (Fig. 5Aa) and their size was measured (Fig. 5Ab). Total LD lipids were extracted and separated by TLC, which shows strain *Bacillus mycoides* contained TAG (Fig. 5Ac), in consistent with the result in Figure 1Dc, lane 5. The total LD proteins were analyzed using MS and the identified proteins were then grouped into 9 categories (Fig. 5Ad). In addition, detailed protein analysis was carried out by slicing the main protein bands and analyzing them using proteomic study (Fig. 5Ae). LDs from strain *Sphingobacterium ginsenosidimutans* of phylum *Bacteroidetes* were also isolated and analyzed as same as *Bacillus mycoides* and strains from other phyla. The results include LD staining (Fig. 5Ba), size measurement (Fig. 5Bb), neutral lipid composition (Fig. 5Bc), proteome (Fig. 5Bd), and main band analysis (Fig. 5Be).

**Figure 5.**
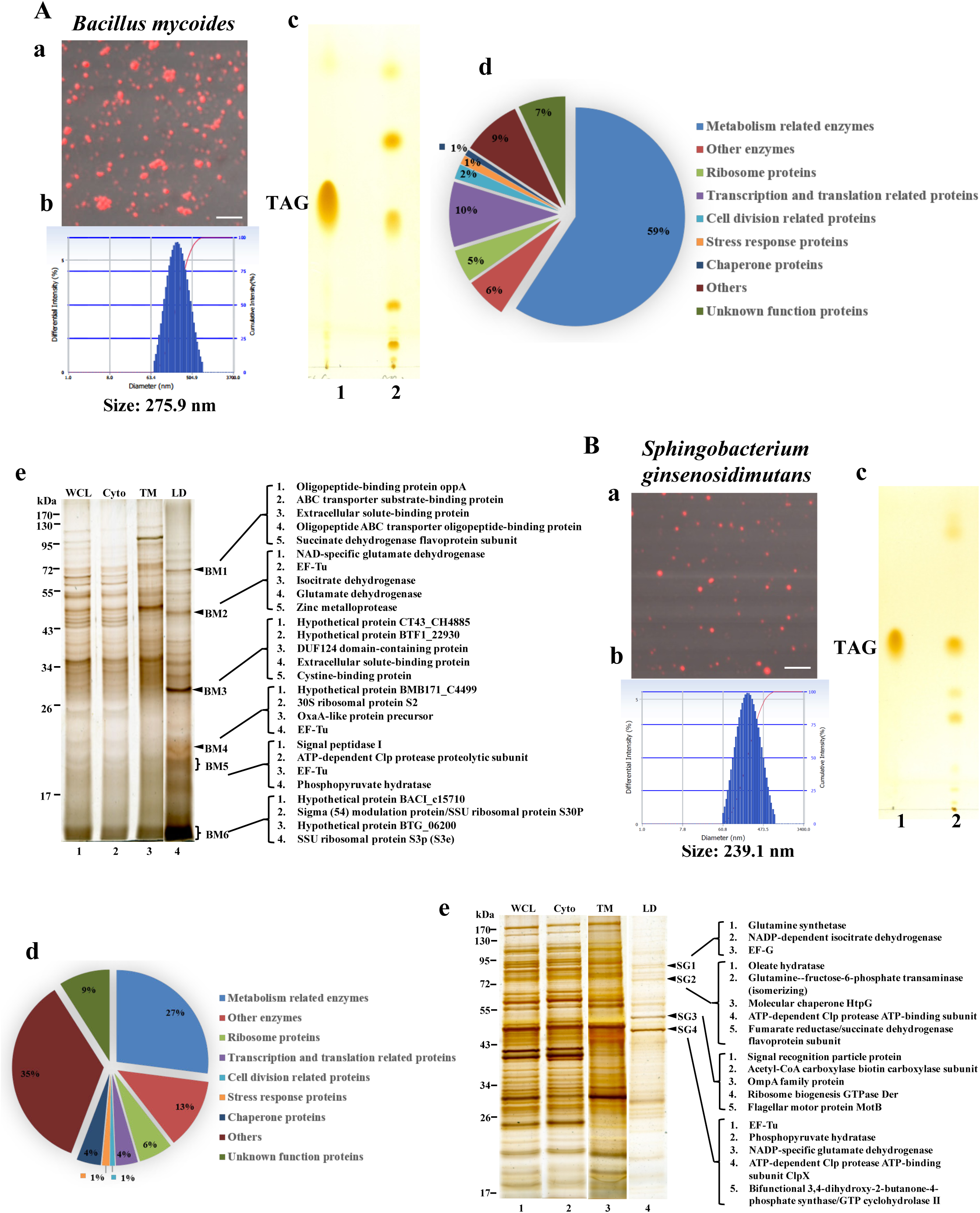
Isolation and Analysis of Lipid Droplets from Phyla *Firmicutes* and *Bacteroidetes*. Bacterial strains were cultivated in the conditions and lipid droplets (LDs) were isolated as described in Materials and Methods. The isolated LDs were stained by LipidTOX Red and imaged using confocal microscopy. Their lipids were extracted, separated and determined using TLC. Their proteins were identified by proteomic studies. **A** LDs were isolated from *Bacillus* of phylum *Firmicutes*. **B** LDs were isolated from *Sphingobacterium* of phylum *Bacteroidetes*. **A** and **B a** The isolated LDs were stained by LipidTOX Red and imaged using confocal microscopy. Scale bar = 2 µm. **b** The size of isolated LDs was measured using Delsa Nano C particle analyzer. **c** The neutral lipids of isolated LDs were analyzed using TLC. **d** The whole proteins from isolated LDs were identified using shotgun mass spectrometry analysis and the identified proteins were categorized by their functions. **e** The whole proteins from isolated LDs (LD) were separated by SDS-PAGE, silver stained, and compared with whole cell lysate (WCL), cytosol (Cyto), and total membrane (TM). Then the major protein bands in LDs were sliced and subjected to mass spectrometry analysis, and identified proteins were marked.

Last, a strain *Nostoc punctiforme* of *Cyanobacteria* was cultured, their LDs were isolated, and LD proteins were analyzed by silver staining. Figure S4R shows that its LD protein composition was unique compared with WCL, Cyto, and TM, suggesting a successful LD isolation.

Based on the accumulated results shown in Figures 1 to 5, especially the isolations and analyses of LDs from many bacterial strains of these five phyla, we finally demonstrate the existence of the organelle in these bacteria, which suggests that LDs are potentially popular in Bacteria domain.

### The lipid droplet is an intrinsic organelle in bacteria

In the above screening results, we have found a number of LD-containing bacteria which are in many phyla. But the LD is not detected in all bacterial strains of these phyla. The representative bacterium is *E. coli*, which is used as a negative control in the above analysis. The original *E. coli* had no LipidTOX Red-positive structures (Fig. S6Aa) and contained very little TAG (Fig. S6Ac). After wax ester synthase/acyl-coenzyme A:diacylglycerol acyltransferase (WS/DGAT) were expressed in the bacterium, LipidTOX Red-positive structures can be detected and a lot of TAG is accumulated in the cells (Fig. S6Aa-c, and [41]). We then purified the possible LDs from the two strains (2119 and 2053) of engineered *E. coli* (Fig. S6Ba) and found that the global structures were stained by LipidTOX Red and the size was also about 200 nm (Fig. 6Aa-c and 6Ba-c). TAG was the major lipid of the purified structures (Fig. 6Ad, 6Bd, and S6Bb). MS results show that LD-associated proteins mainly contained metabolism-related enzymes and other enzymes, ribosome proteins, and DNA-binding proteins (Fig. 6Ae and 6Be), which was similar to other bacterial LD proteomes. Meanwhile, the major proteins were also revealed and PspA was one of the major LD proteins (Fig. 6Af, 6Bf and S6Bc). The data demonstrate that the engineered *E. coli* contain LDs, which suggests that they possess other components of LD formation except for TAG synthesis. These results indicate that the bacteria in which the LD is not detected may have the ability to form LDs if the lost components are put back. Furthermore, we found that not only soil bacteria contained LDs, LDs were also detected in some human symbiotic bacteria including *Bifidobacterium longum* (anaerobe, Fig. S4Aa-c), *Acinetobacter calcoaceticus* (aerobe, Fig. S4Ma and b), and *Mycobacterium smegmatis* (aerobe, Fig. S5). Altogether, these results suggest that the LD is an intrinsic organelle in bacteria in wild or symbiotic environment, with or without oxygen, even though it may be lost based on the bacterial needs.

**Figure 6.**
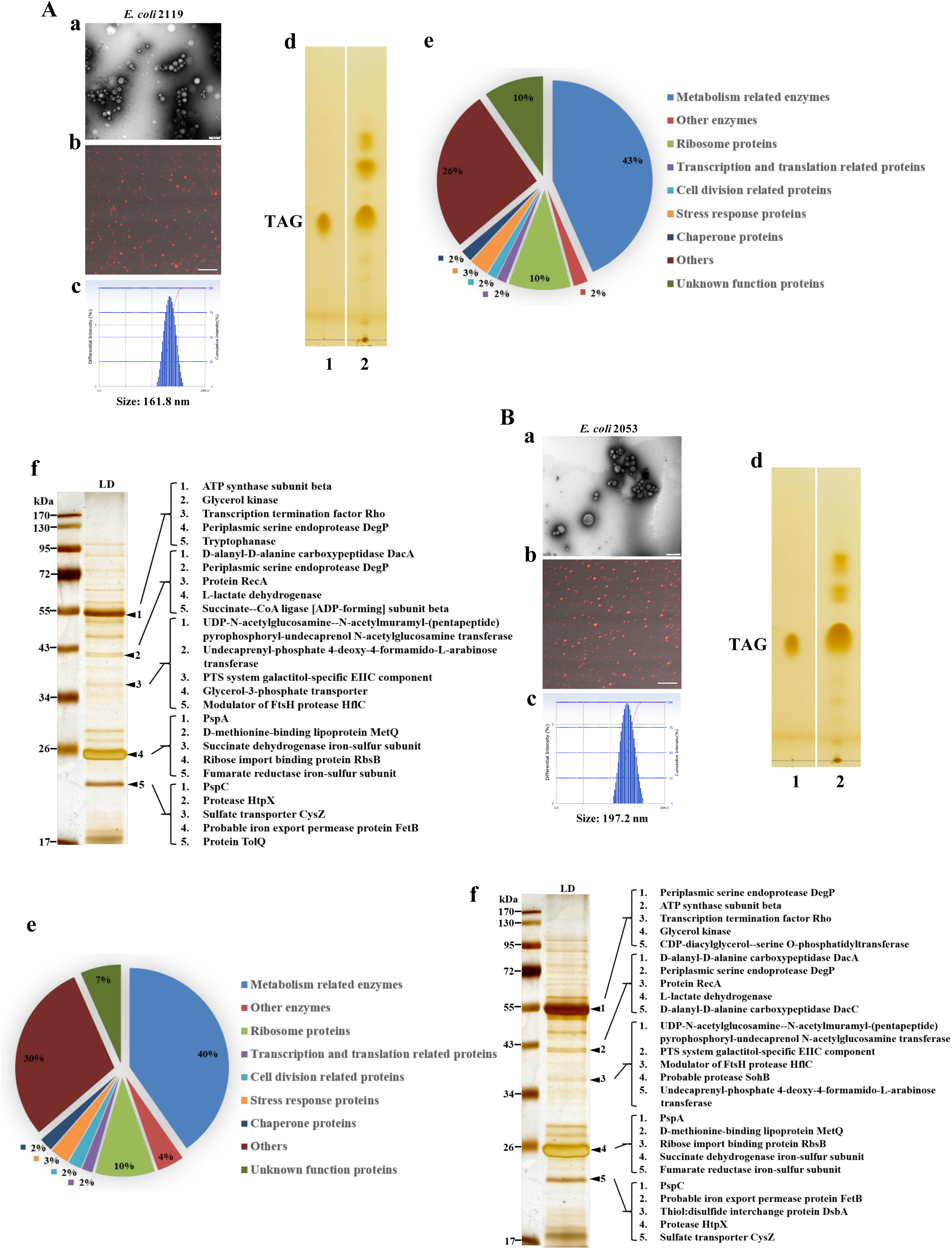
Isolation and Analysis of Lipid Droplets from Engineered *E. coli*. Bacterial strains were cultivated in the conditions and lipid droplets (LDs) were isolated as described in Materials and Methods. The isolated LDs were stained by LipidTOX Red and imaged using confocal microscopy. Their lipids were extracted, separated and determined using TLC. Their proteins were identified by proteomic studies. **A** and **B** Isolated LDs from Engineered *E. coli* 2119 and 2053, respectively, imaged by TEM after negative staining (**a**) (scale bar = 500 nm), confocal microscopy (**b**) (scale bar = 2 µm), and size measurement (**c**). **d** The neutral lipids of isolated LDs were analyzed using TLC. **e** The whole proteins from isolated LDs were identified using shotgun mass spectrometry analysis and the identified proteins were categorized by their functions. **f** The whole proteins from isolated LDs (LD) were separated by SDS-PAGE, silver stained. Then the major protein bands in LDs were sliced and subjected to mass spectrometry analysis, and identified proteins were marked.

### The lipid droplets are conserved and inheritable in Bacteria domain

The existence of LDs in many bacterial strains of five major phyla indicates that LDs in different bacteria may be relevant. Therefore, we then conducted some experiments to test whether LD is a conserved organelle in Bacteria domain. LDs mainly contain lipids and proteins. As shown in the above results, although their major neutral lipids were similar in some bacteria and distinct in others, they were all stained by neutral lipid dye LipidTOX, had sphere-shaped morphology, contained small amount of phospholipids, and had average diameter of about 200 nm. We then focused on proteins to determine if some of LD main proteins are conserved. Based on proteomic analyses of isolated bacterial LDs in this study and a couple of previous reports, it was found that they basically had six functional protein categories, including proteins related to metabolism, DNA binding, transcription, translation, protein folding, and stress response (Fig. 3Dd, 4Bd, 5Ad, 5Bd, 6Ae, 6Be, S5Ae, and S5Be). Therefore, we here selected a stress response protein PspA. In our previous LD proteomics, it was the first time to identify PspA as one of LD major proteins, which was further verified by gene deletion in bacteria *Rhodococcus jostii* RHA1 and *Rhodococcus opacus* PD630 [19, 35]. Its GFP-fusion protein is indeed associated with LDs in *Mycobacterium smegmatis* but lack of ring structure [20]. To further determine the targeting of PspA on LDs, a PspA-GFP fusion protein was expressed in *Rhodococcus jostii* RHA1 and co-localization with LipidTOX-labeled LDs was imaged. We also found the association of PspA with LD but without ring structure either (Fig. S7A, lower panel). PspA is a very widespread protein in Bacteria domain. It was found that it exists in almost all prokaryotes and even in some eukaryotes (Fig. S7B). In the above proteomic analyses, we found that PspA was associated with LDs not only in the *Rhodococcus* (Fig. 3I), *Mycobacterium* (Fig. S5Bd) and *Streptomyces* [49] of the phylum *Actinobacteria*, but also in the *Bacillus* (Table S5) of the phylum *Firmicutes.* Even we also found that the PspA was in the *E. coli* LD proteome (Fig. 6Af and 6Bf), which suggests that PspA from *E. coli* still has its ability of localization on LD. The intrinsic ability of PspA targeting to LDs may also explain why PspA from *E. coli* can localize on the LDs in *Mycobacterium smegmatis* [50]. These results suggest that PspA is not only a widespread protein existing in Bacteria domain, but also a universal LD protein in bacteria. Altogether, these findings further verified that the organelle LD is conserved in Bacteria domain.

Together these data and findings suggest that the LD is conserved in Bacteria domain. So, whether it can be inherited from mother cell is our next aim to study. To answer the question, we used *Rhodococcus jostii* RHA1 as model bacterium since it can form branches when it grows and divides [51], which can allow us to observe LD dynamic during bacterial division. The bacteria were observed via time-lapse confocal microscopy and LDs were stained by LipidTOX Red (Fig. 7A). It was revealed that along with the birth and growth of bacterial branches, LDs were inherited from a mother cell into daughter cells (Fig. 7A). Then we performed 3D tomography at bacterial branching site and found that several LDs were dividing at the sites (Fig. 7B). Moreover, the TEM image confirms the bacterial branching and LD inheritance (Fig. S8A). Altogether, these data suggest that the LD can be inherited from a mother cell to daughter cells.

**Figure 7.**
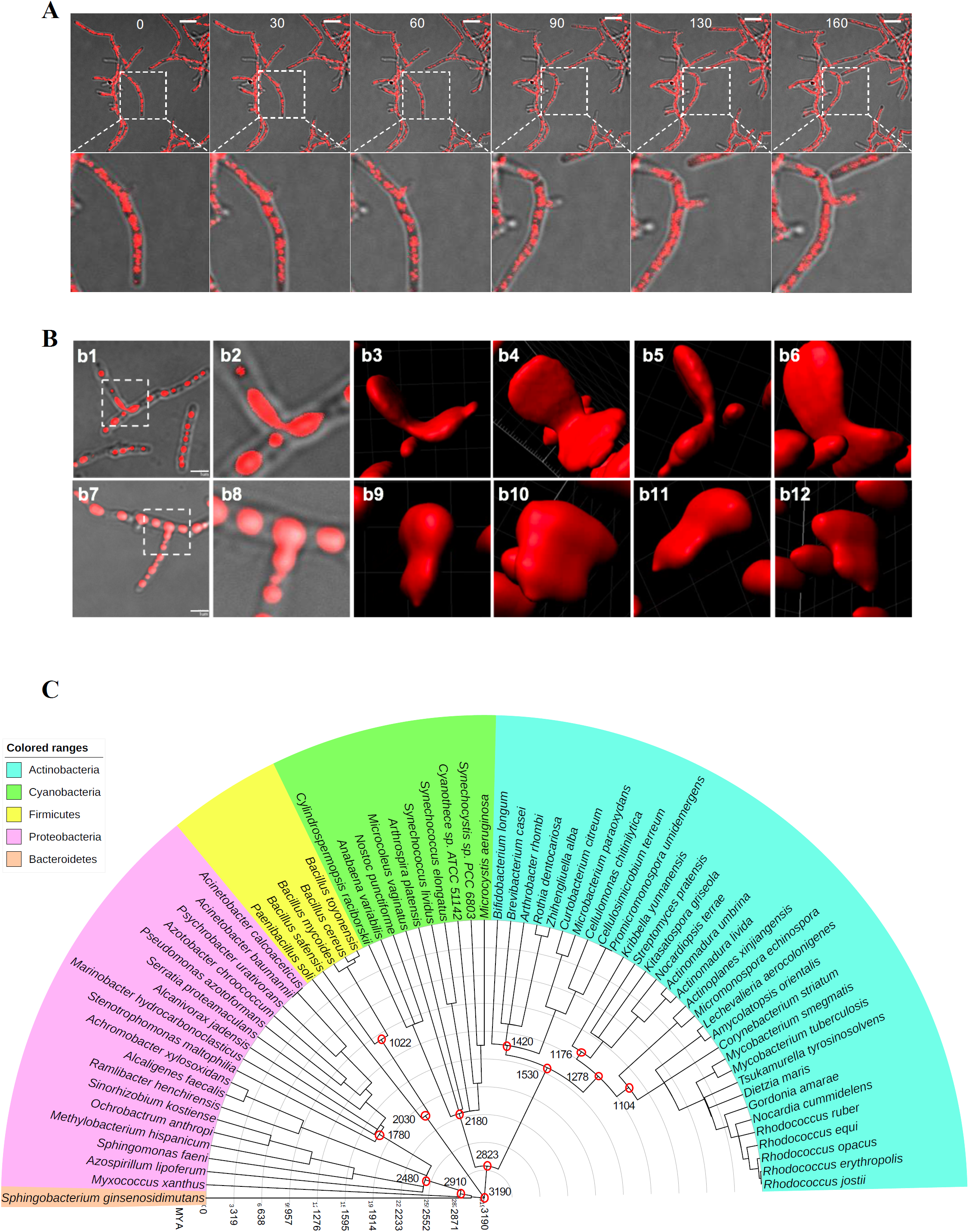
The Lipid Droplet Is an Ancient and Inheritable Organelle in Bacteria Domain. **A** The time-lapse live imaging of the bacterium *Rhodococcus jostii* RHA1. The cell growth and division of one bacterium as well as the inheritance of the LDs were observed. The numbers indicate the minutes. The bacteria were stained by LipidTOX Red and observed by confocal microscope. The Scale bar = 5 μm. **B** The representatives of the LD inheritance during cell division in the bacterium *Rhodococcus jostii* RHA1. The bacteria were stained by LipidTOX Red and observed by confocal microscope. The images were 3D reconstructed by Imaris software (b3-b6 and b9-b12 with different angles). Scale bar = 2 μm. **C** The phylogenetic analysis of LD-containing bacteria using Timetree database. And the tree was displayed via Interactive Tree Of Life. Cyan, green, yellow, pink, and brown represent *Actinobacteria, Cyanobacteria, Firmicutes, Proteobacteria* and *Bacteroidetes*, respectively. Values in black show the inferred ages of nodes in years MYA (million years ago).

### The lipid droplet is an ancient organelle

The above results have revealed that the LD is widespread in five major bacterial phyla, conserved, and inheritable in Bacteria domain. Furthermore, the previous studies also report that several prokaryotes contain LDs, including other cyanobacteria [52-54] and others [18, 39, 55-58]. All these data indicate LD may be ancient. Thus, we combined previously reported bacteria that had been demonstrated to contain LDs with our finding in this study and performed the phylogenetic analysis on the bacteria containing LDs (Fig. 7C and S8B). According to the estimated time of each node in the evolutionary tree, it shows that the ancestor of these LD-containing bacteria may appear about 3.19 billion years ago (Fig. 7C). The result indicates that the LD may be one of the most ancient cellular membrane-bound organelles in the life evolution.

## Discussion

Lipid droplet (LD) has a monolayer phospholipid membrane that is unique compared to other cellular structures with a bilayer phospholipid membrane. The property of this unique membrane provides a specific environment for certain membrane proteins. Accumulated evidence demonstrates that some membrane proteins are only localized on LDs. These proteins are degraded by the proteasome during LD lipolysis. We term them LD resident proteins, such as PLIN1, ADRP/PLIN2, DHS-3, MDT-28, and MLDS [6]. Interestingly, LD membrane also contain most features of bilayer membrane. Except transmembrane proteins, most peripheral membrane proteins can also be localized and function on LDs. For example, some membrane trafficking proteins are found to be associated with LDs [11] such as Rabs and SNAREs, and plasma membrane protein caveolin1/2 have also been identified on LDs, demonstrating that the LD monolayer membrane can function as bilayer phospholipid membranes.

The monolayer membrane not only covers and protects neutral lipids but also provides a distinct solid face that is different from bacterial plasma membrane. Therefore, the LD in bacteria supplies a unique membrane structure for a specialized compartment. On the one hand, the LD stores energy and carbon source. On the other hand, the LD presents a unique solid surface that differs from host plasma membrane for some biological processes. For instance, we recently found that bacterial LDs bind genomic DNA and protect DNA from damage in extreme living conditions [17]. These facts suggest that phospholipid monolayer structure, LD, was generated to form a unique membrane compartment that not only stores energy to increase metabolic efficiency, but also protects genomic DNA to facilitate heritage. Therefore, bacterial LD meets the most basic necessities of living organisms, metabolism and heritage, which indicates the evolutionary significance of the LD as membrane-bound structure in early cellular life.

In addition, our study presented that at least three major neutral lipids TAG, WE, and PHA/B were detected in these bacterial LDs (Fig. 3Bc, 3Cc, 3Dc, 4Bc, 5Ac, 5Bc, 6Ad, 6Bd, S4Ac, S4Bc, S4Cc, S4Dc, S5Ad, S5Bd, S5Cd, and S6Bb), which is similar to eukaryotic LDs that mainly contain TAG, CE/SE, ether lipids, and RE [29, 30]. The content of LDs, in many cases, is not simply for energy storage. In fact, LDs store many precursors for the metabolites that living organisms use. Therefore, LD provides a special compartment, reserves energy and carbon source, and stores biological building blocks and precursors.

Although our current study shows existence of LDs in many types of bacteria, but in some bacteria, LDs could not be detected or were very rare (Fig. 1B). It is possible that these bacteria were not cultured in proper conditions or they lost certain essential genes for accumulating LDs. For example, the *E. coli* TOP10 had very low level of TAG (Fig. S6). But after the WS/DGAT was expressed in the bacterium, we identified LDs from the engineered bacteria (Fig. 6). The proteomic study revealed that PspA was also one of the LD major proteins same as other bacteria, suggesting that the *E coli* TOP10 has the ability to form LDs and the LDs are conserved in Bacteria domain. Additionally, a recent study reports that PspA targets bacterial LDs using an amphipathic α-helix [50], which is similar with the pattern of mammalian LD protein localization [6]. Furthermore, current study also found that PspA was conserved from bacteria, archaea, to eukaryotes (Fig. S7B). These results further support the hypotheses that most of bacteria may have either LDs or ability to generate LDs, a membrane-bound organelle, and LDs are conserved from bacteria to humans [28].

Although membrane-bound organelles are essential for eukaryotic cells, their origin or/and initiation is still elusive. Except mitochondria and chloroplasts in eukaryotes that may be evolved through endosymbiosis from certain bacterial invasion, existence of membrane-bound organelles and where they come from are still a mystery. Based on previous studies, a few of membrane-like structures have been identified in several bacterial strains [59], including membrane-like structure in *Cyanobacteria* and *methanotrophic* bacteria [60]. Other potential membrane structures include anammoxosome [61], magnetosome [62], acidocalcisome [63], chlorosomes [64], and pirellulosomes [65]. These structures are rare, very conditional, and some may not be independent organelles or intracellular membrane structures. In contrast, LDs were well distributed, conserved, and independent organelles in bacteria and may be evolved earlier than other.

According to Charles Darwin’s theory of universal common ancestry and the current knowledge, the most recent common ancestor of all cellular life on Earth is called the last universal common ancestor (LUCA). A recent study using a molecular clock model suggests that the LUCA existed 3.9 billion years ago (the end of The Late Heavy Bombardment) [66]. In this study, we collected and analyzed the bacteria from normal environmental soil and human body as well as engineered bacteria, and then performed phylogenetic analysis of the representative strains of LD-containing bacteria. Based on the above analyses, it was found that the bacteria with LDs can be predated to 3.19 billion years ago (Fig. 7C). Here we name the ancestor with the LD as the Last Universal Common Ancestor with Lipid Droplet (LUCALD). Since some studies report that the LD-like structures are also found in archaea [1, 38], it is possible that the emergence of LUCALD may be earlier than 3.19 billion years. Moreover, LDs were found in anaerobic bacteria (Fig. S4Aa-c), suggesting that oxygen is not necessary for the LD emergence, which supports that LDs may exist before atmospheric oxygen rising (Proterozoic). Furthermore, the time-lapse imaging and 3D tomography reveal that LDs are inherited during bacterial division (Fig. 7A and 7B), which is similar to LDs in eukaryotes [67-70]. The LD inheritance further supports the hypothesis that the LD in bacteria and eukaryotes has the common ancestor. Altogether, the analyses in this study indicate that the LD may be the first membrane-bound organelle in all cellular life on Earth and conserved to humans.

## Acknowledgements

We would like to thank Ms. Zhensheng Xie for her technical supports of proteomic study, Mrs. Yan Teng (Center for Biological Imaging, IBP, CAS) for her help of taking and analyzing Confocal images, Ms. Shuoguo Li (Center for Biological Imaging, IBP, CAS) and Ms. Yun Feng (Center for Biological Imaging, IBP, CAS) for their help of taking Imaris analysis. This work was supported by the Natural Science Foundation of China (Grant No. 91857201), National Key R&D Program of China (Grant No. 2016YFA0500100, 2018YFA0800900 and 2018YFA0800700), and National Natural Science Foundation of China (Grant No. 31671402, 91954108, 31671233, 31771314, 31701018 and U1702288). This work was also supported by the “Personalized Medicines——Molecular Signature-based Drug Discovery and Development”, Strategic Priority Research Program of the Chinese Academy of Sciences, Grant No. XDA12040218, and China Postdoctoral Science Foundation (BX20180345).

## Author contributions

P.L. and C.Z. conceived the project and designed the experiments. X.C., O.O.O., and C.Z. performed the most experiments. Z.L., X.L., and M.Z. assisted to carry out experiments, F.S., X.W., and L.Z. provided the bacterial source. Z.Z., J.W., M.A.H., and C.Z. contributed to the phylogenetic analyses. X.C. and S.Z. conducted electron microscopy observation. Experiments and manuscript were assisted by contributions from X.Z., X.W., L.Z. P.L., C.Z., X.C., and O.O.O. organized the Data. Manuscript was written by C.Z. and P.L. All authors have read and approved the manuscript.

