## Supplemental Figure Legends for "Lipid Droplet Is an Ancient and Inheritable Organelle in Bacteria"

**Figure S1 Screening of Lipid Droplet-contained Bacteria**

**A** *Rhodococcus jostii* RHA1 was cultivated in the conditions described in Materials and Methods. **a** Bacteria were visualized using DIC. **b** Neutral lipids in these strains were stained by LipidTOX red and imaged using confocal microscopy. **c** Lipid droplet (LD)-like structures were visualized using transmission electron microscopy (TEM). **B** Procedure for the screening of lipid droplet-contained soil bacteria.

**Figure S2 Screening of Neutral Lipid-accumulating Bacteria by TLC**

Bacterial strains were cultivated in the conditions and lipid droplets (LDs) were isolated as described in Materials and Methods. Total lipids of the isolated LDs were extracted using Bligh-Dyer method, and neutral lipids were separated and determined using TLC. **A** Bacterial strains of phylum *Actinobacteria*. **B** Bacterial strains of phylum *Proteobacteria*. **C** Bacterial strains of phylum *Firmicutes*. **D** Bacterial strains of phylum *Bacteroidetes*.

**Figure S3 Morphological Screening of Neutral Lipid-contained Bacteria**

Bacterial strains were cultivated in the conditions described in Materials and Methods. Neutral lipids in these strains were stained by LipidTOX red and imaged using confocal microscopy. Scale bar=5 µm. **A** The images of neutral lipids in strains of phylum *Actinobacteria*. **B** The images of neutral lipids in strains of phylum *Proteobacteria*. **C** The images of neutral lipids in strains of phylum *Firmicutes*. **D** The image of neutral lipids in strains of phylum *Bacteroidetes*. **E** The image of neutral lipids in strains of phylum Cyanobacteria.

**Figure S4 Isolation and Analysis of Lipid Droplets from Bacteria**

Bacterial strains were cultivated in the conditions and lipid droplets (LDs) were isolated as described in Materials and Methods. The isolated LDs were stained by LipidTOX red and imaged using confocal microscopy. Their lipids were extracted using Bligh-Dyer method, and neutral lipids were separated and determined using TLC. **A** to **D** **a** The isolated LDs were stained by LipidTOX red and imaged using confocal microscopy. Scale bar=2 µm. **b** The size of isolated LDs was measured using Delsa Nano C particle analyzer. **c** The neutral lipids of isolated LDs were analyzed using TLC*.* **A** LDs were isolated from *Bifidobacterium* of phylum *Actinobacteria*. **B** LDs were isolated from *Streptomyces* of phylum *Actinobacteria*. **C** LDs were isolated from *Kitasatospora* of phylum *Actinobacteria*. **D** LDs were isolated from *Tsukamurella* of phylum *Actinobacteria*. **E** and **F** **a** The isolated LDs were stained by LipidTOX red and imaged using confocal microscopy. Scale bar=2 µm. **b** The size of isolated LDs was measured using Delsa Nano C particle analyzer. **E** LDs were isolated from *Pseudomonas* of phylum *Proteobacteria*. **F** LDs were isolated from *Ochrobacterium* of phylum *Proteobacteria*. **G** **a** The isolated LDs were stained by LipidTOX red and imaged using confocal microscopy. Scale bar=2 µm. **b** The size of isolated LDs was measured using Delsa Nano C particle analyzer. **c** The whole proteins from isolated LDs (LD) were separated by SDS-PAGE, stained by silver staining, and compared with whole cell lysate (WCL), cytosol (Cyto), and total membrane (TM). LDs were isolated from *Amycolatopsis orientalis* of phylum *Actinobacteria*. **H** to **M** **a** The isolated LDs were stained by LipidTOX red and imaged using confocal microscopy. Scale bar=2 µm. **b** The whole proteins from isolated LDs (LD) were separated by SDS-PAGE, stained by silver staining, and compared with whole cell lysate (WCL), cytosol (Cyto), and total membrane (TM). **H** LDs were isolated from *Rhodococcus* sp*.* of phylum *Actinobacteria*. **I** LDs were isolated from *Streptomyces tanashiensis* of phylum *Actinobacteria*. **J** LDs were isolated from *Streptomyces mutomycini* of phylum *Actinobacteria*. **K** LDs were isolated from *Curtobacterium citreum* of phylum *Actinobacteria*. **L** LDs were isolated from *Streptomyces griseus* of phylum *Actinobacteria*. **M** LDs were isolated from *Acinetobacter calcoaceticus* of phylum *Proteobacteria*. **N** to **R** The whole proteins from isolated LDs (LD) were separated by SDS-PAGE, stained by silver staining, and compared with whole cell lysate (WCL), cytosol (Cyto), and total membrane (TM). **N** LDs were isolated from *Brevibacterium casei* of phylum *Actinobacteria*. LDs were isolated from *Microbacterium esteraromaticum* of phylum *Actinobacteria*. **O** LDs were isolated from *Acinetobacter baumannii* of phylum *Proteobacteria* . **P** LDs were isolated from *Arthrobacter rhombi* of phylum *Actinobacteria*. **Q** LDs were isolated from *Actinomycete* HVG71 of phylum *Actinobacteria*. **R** LDs were isolated from *Nostoc punctiforme* of phylum *Cyanobacteria*.

**Figure S5 Isolation and Proteomic Analysis of Lipid Droplets from Bacteria**

Bacterial strains were cultivated in the conditions and lipid droplets (LDs) were isolated as described in Materials and Methods. The isolated LDs were stained by LipidTOX red and imaged using confocal microscopy. Their lipids were extracted using Bligh-Dyer method, and neutral lipids were separated and determined using TLC. Their proteins were identified by proteomic studies. **A a** The isolated LDs from *Mycobaterium smegmatis* of phylum *Actinobacteria* were stained by LipidTOX Red and imaged using confocal microscopy. **b** The isolated LDs were visualized by TEM. Scale bar=2 µm. **c** The size of isolated LDs was measured using Delsa Nano C particle analyzer. **d** The neutral lipids of isolated LDs were analyzed using TLC*.* **e** The whole proteins from isolated LDs (LD) were separated by SDS-PAGE, stained by silver staining, and compared with whole cell lysate (WCL), cytosol (Cyto), and total membrane (TM). Then the major protein bands in LDs were sliced and subjected to mass spectrometry analysis, and identified proteins were marked. **f** The whole proteins from isolated LDs were identified using shotgun mass spectrometry analysis and the identified proteins were categorized by their functions. **B** and **C** **a** The bacteria were visualized using DIC and neutral lipid staining (**a** Biodipy and **b** LipidTOX). **b** The isolated LDs were stained by LipidTOX Red and imaged using confocal microscopy. Scale bar=2 µm. **c** The size of isolated LDs was measured using Delsa Nano C particle analyzer. **d** The neutral lipids of isolated LDs were analyzed using TLC*.* **e** The whole proteins from isolated LDs (LD) were separated by SDS-PAGE, stained by silver staining, and compared with whole cell lysate (WCL), cytosol (Cyto), and total membrane (TM). Then the major protein bands in LDs were sliced and subjected to mass spectrometry analysis, and identified proteins were marked. **f** The whole proteins from isolated LDs were identified using shotgun mass spectrometry analysis and the identified proteins were categorized by their functions. **B** LDs were isolated from *Sinorhizobium* sp. of phylum *Proteobacteria*. **C** LDs were isolated from *Stenotrophomonas maltophilia* of phylum *Proteobacteria*.

**Figure S6 Generation of Lipid Droplets by Engineered *E. coli***

Bacterial strains were cultivated in the conditions described in Materials and Methods. Neutral lipids in these strains were stained by LipidTOX red and imaged using confocal microscopy. **A a** Confocal microscopy imaging of neutral lipids in engineered *E. coli*. The commercial *E. coli* strain BL21 was used as negative control. Scale bar=2 µm. **b** The ultra-structure of the representative strains was observed using TEM. Briefly, the cells were dehydrated through an ethanol series after a two-step fixation. The samples were then embedded in resin and thin sectioned. The sections were stained and visualized by the TEM. **c** Neutral lipids were separated and determined by thin layer chromatography (TLC). Lane 1-3 are *E. coli* BL21, engineered *E. coli* 2053 and 2119 respectively. **B a** Engineered *E. coli* cells cultivated in LB were collected and homogenized. After removal of cell debris, whole cell lysate ultracentrifugation was performed at 182,000 g for 1 h at 4°C (Beckman SW40). The lipid droplet fraction floated to the top of the sucrose gradient. **b** The neutral lipids of isolated LDs were analyzed using TLC. **c** LD proteins were extracted from engineered *E. coli* and separated by 10% SDS-PAGE followed by silver staining.

**Figure S7 PspA Is a Conserved Protein on Bacterial Lipid Droplets**

**A** The localization of PspA fused with GFP in the bacterium *Rhodococcus jostii* RHA1. The bacteria were stained by LipidTOX red and observed by confocal microscope. The Scale bar = 5 μm. **B** The evolutionary analysis of PspA modified from Pfam database.

**Figure S8 Lipid Droplet Is an Ancient and Inheritable Organelle**

**A** The ultra-structure of the *Rhodococcus jostii* RHA1 was observed using TEM. The enlarged image is on the right. Scale bar = 500 nm. **B** The phylogenetic analysis of bacteria containing LDs in this study by 16S rRNA maximum likelihood method.
