## Supplemental Figures for "Lipid Droplet Is an Ancient and Inheritable Organelle in Bacteria"

### Figure S1 Screening of Lipid Droplet-contained Bacteria

A

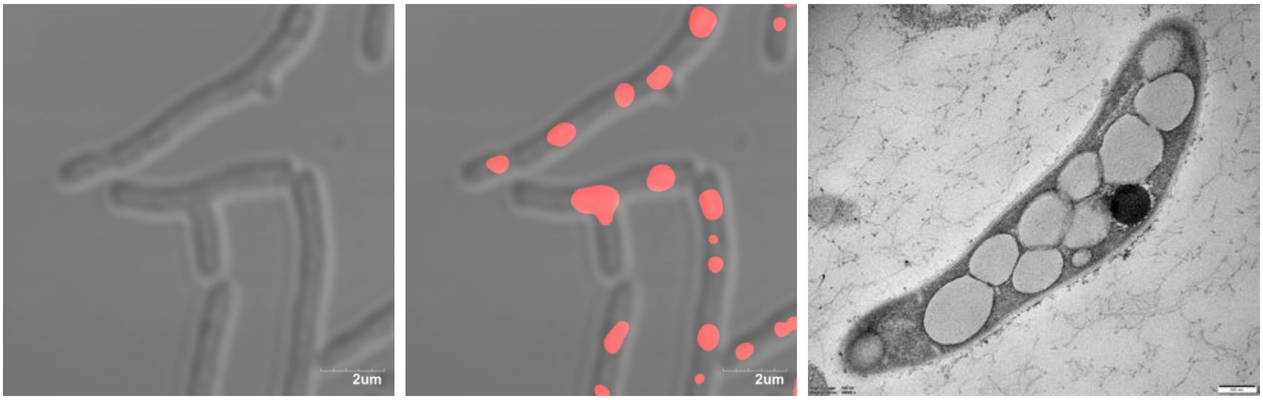


B

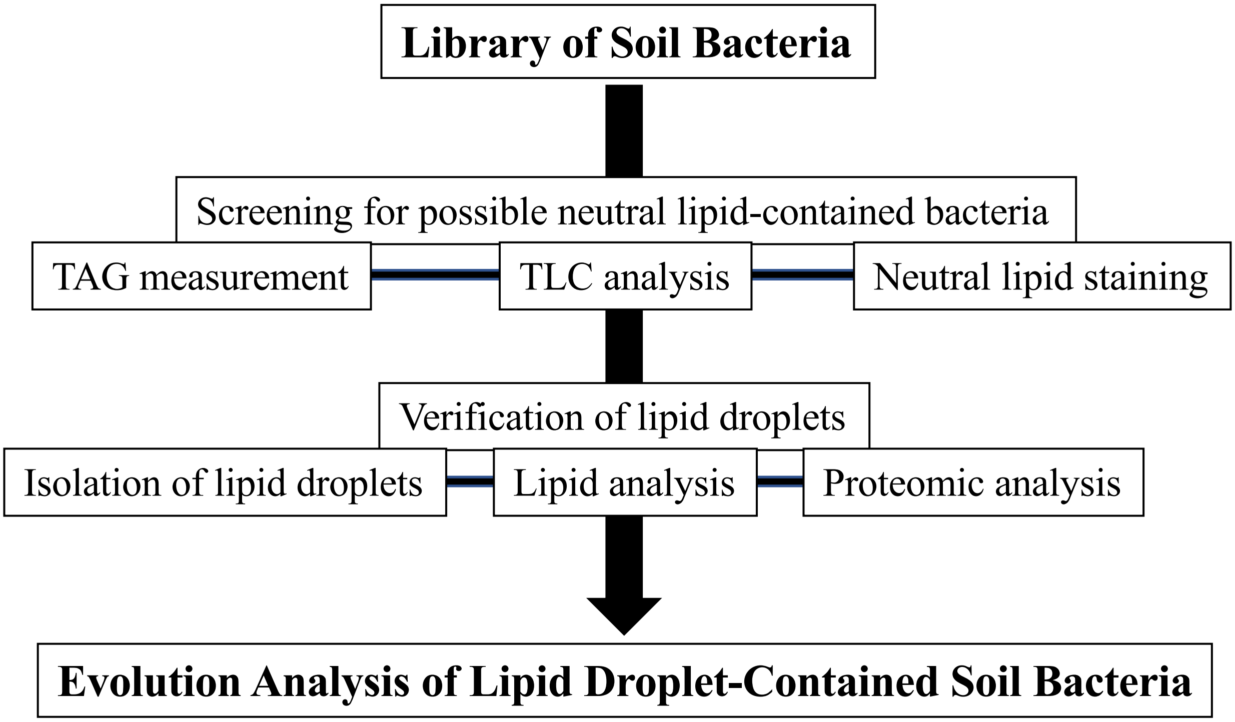

### Figure S2 Screening of Neutral Lipid-accumulating Bacteria by TLC

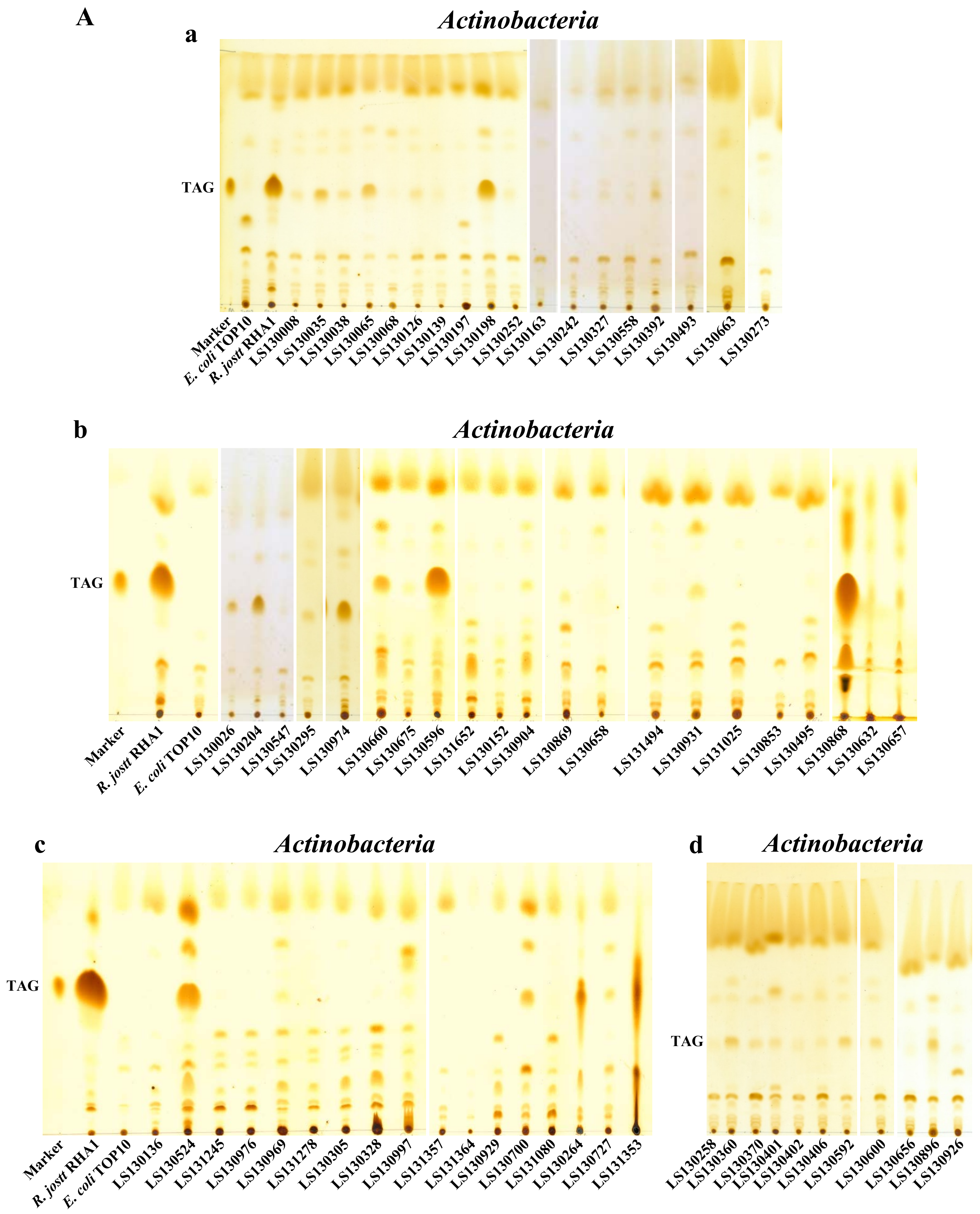

### Figure S2 Screening of Neutral Lipid-accumulating Bacteria by TLC

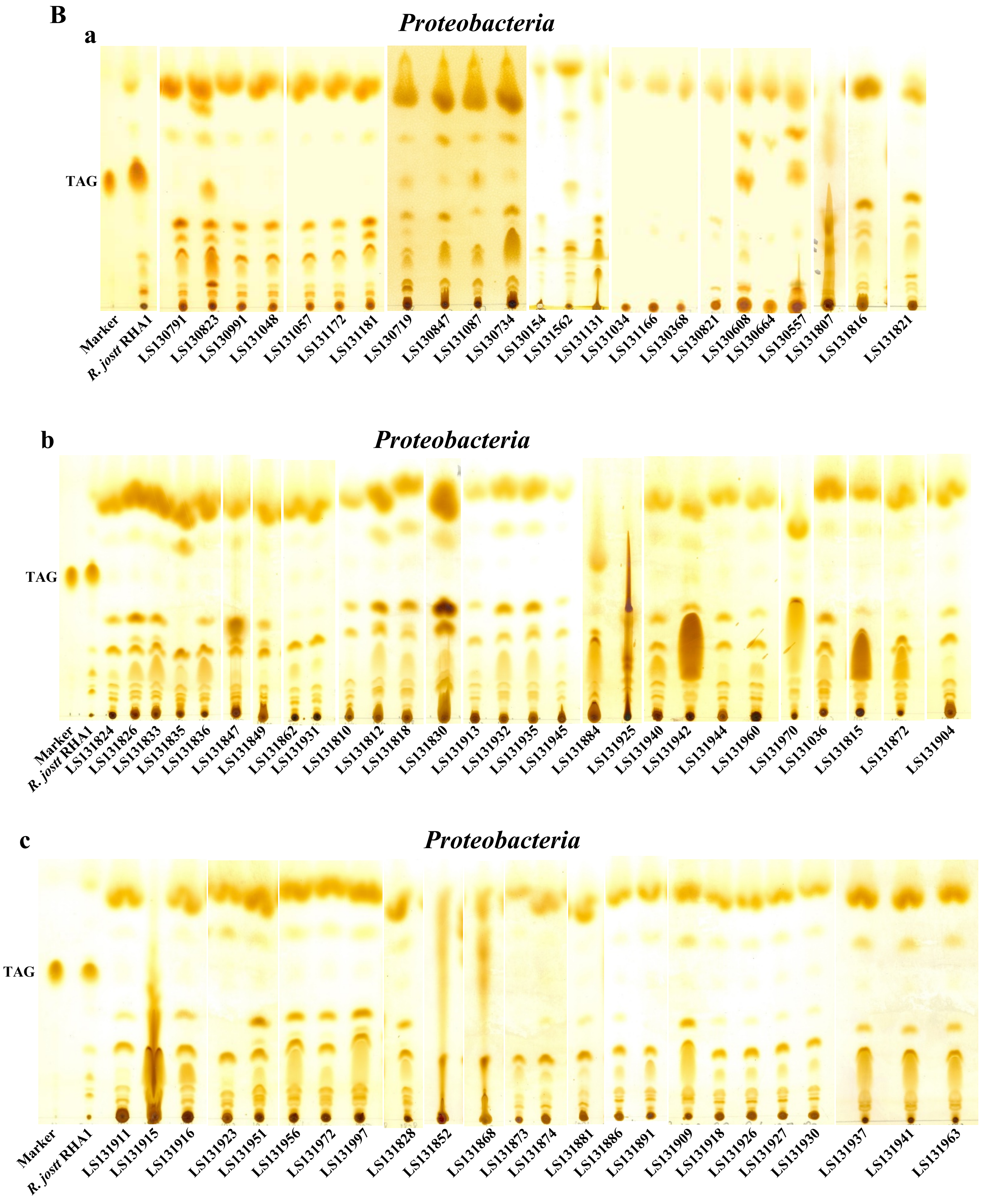

### Figure S2 Screening of Neutral Lipid-accumulating Bacteria by TLC

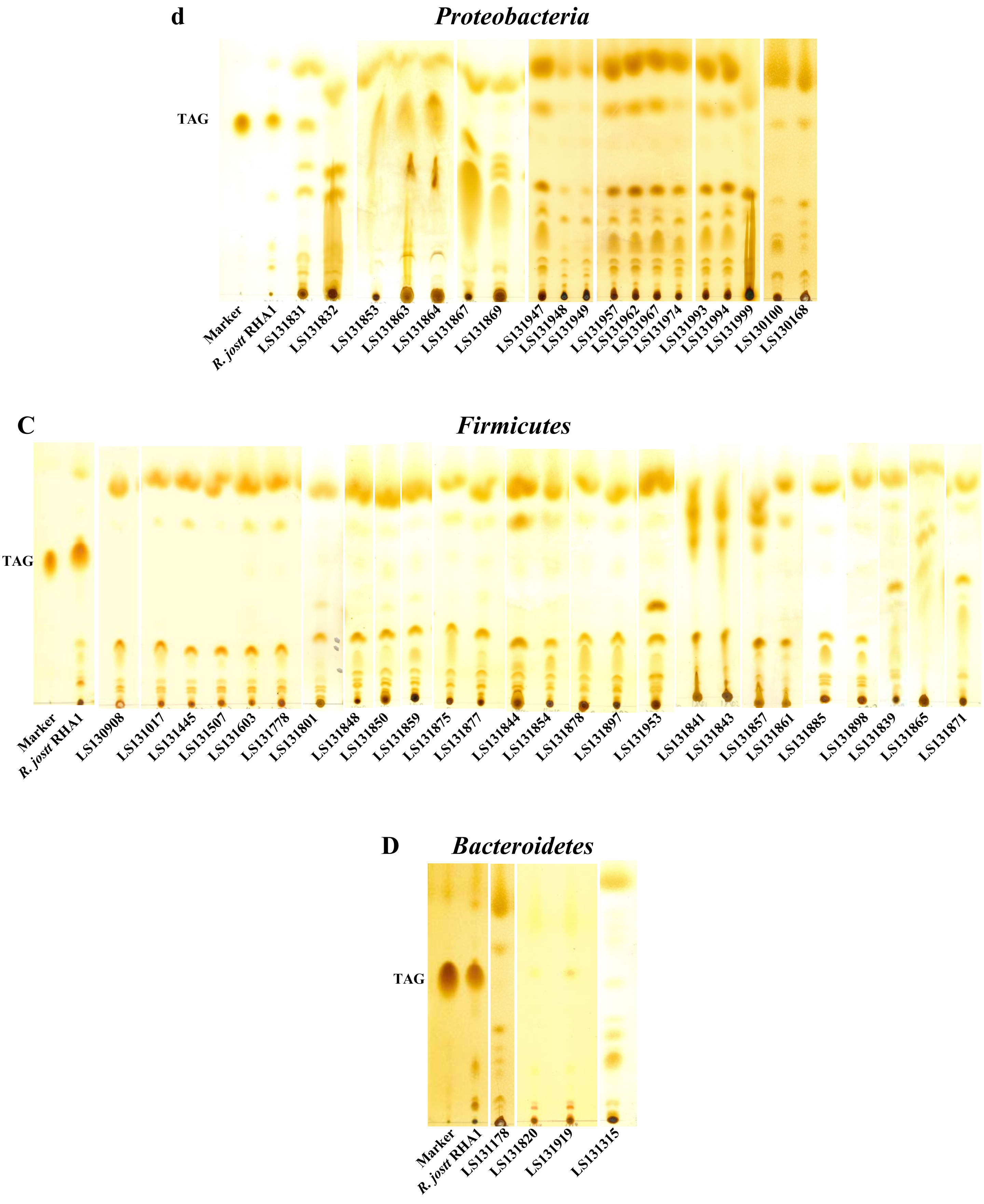

### Figure S3 Morphological Screening of Neutral Lipid-contained Bacteria

**A**

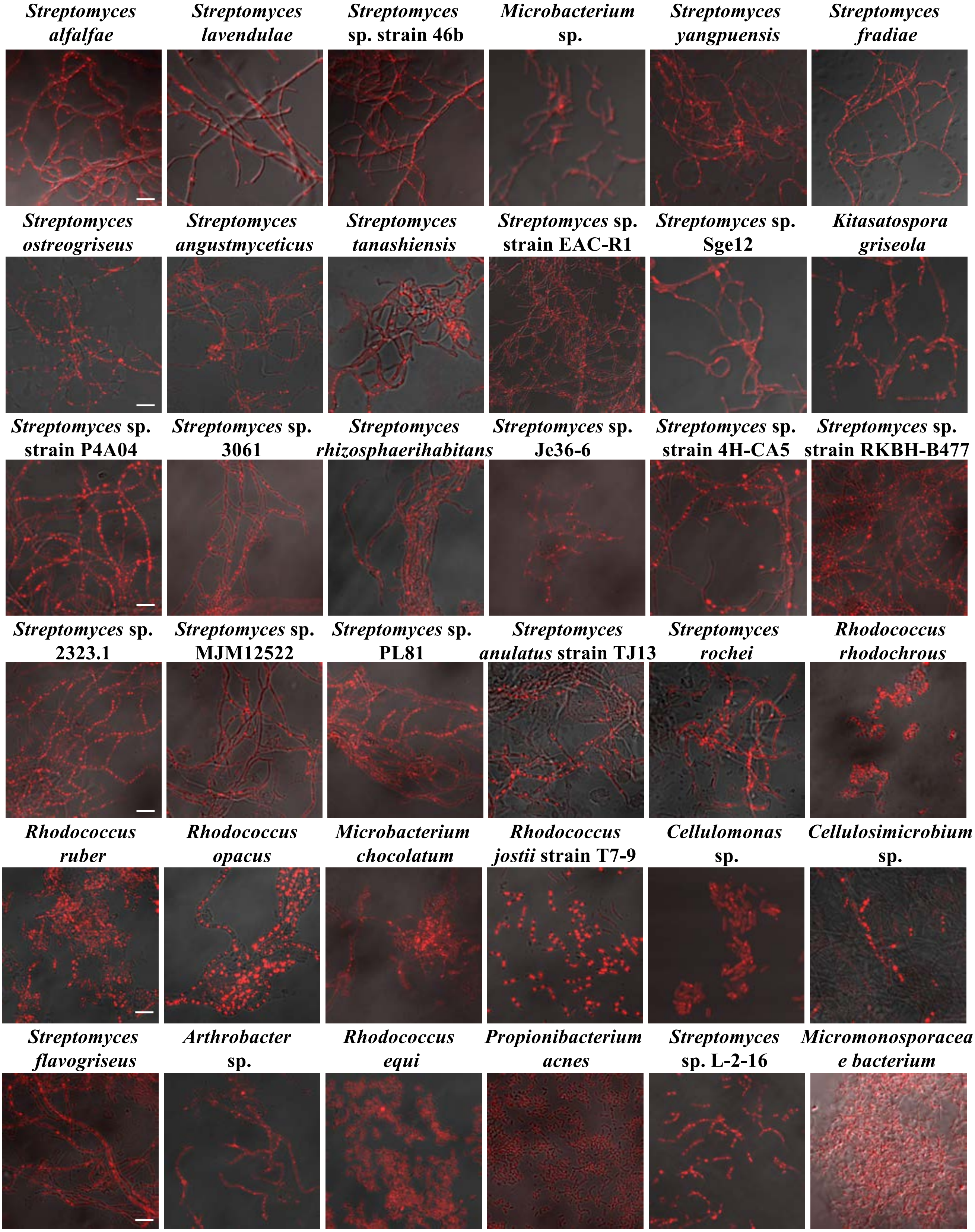

### Figure S3 Morphological Screening of Neutral Lipid-contained Bacteria

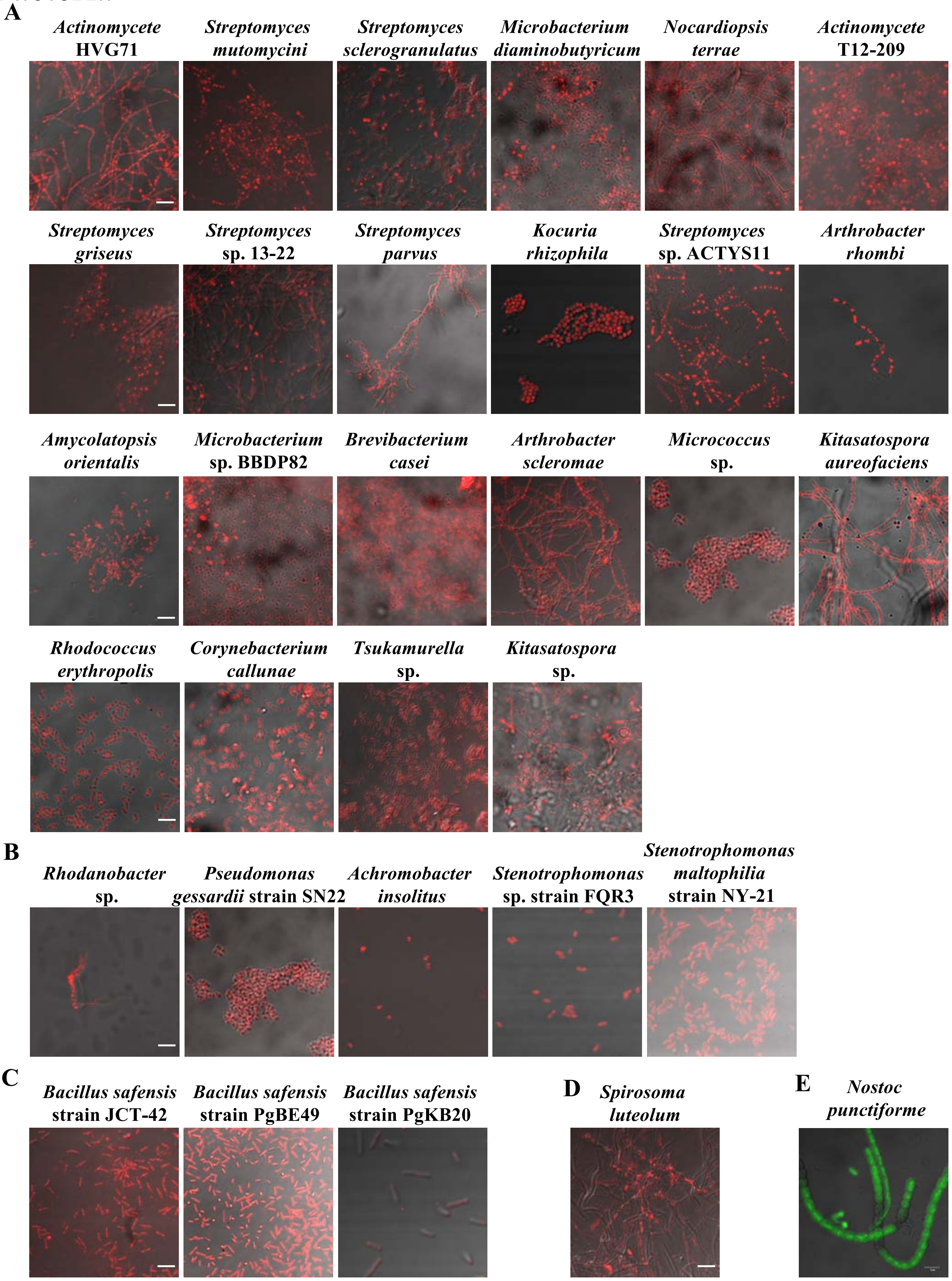

### Figure S4 Isolation and Analysis of Lipid Droplets from Bacteria

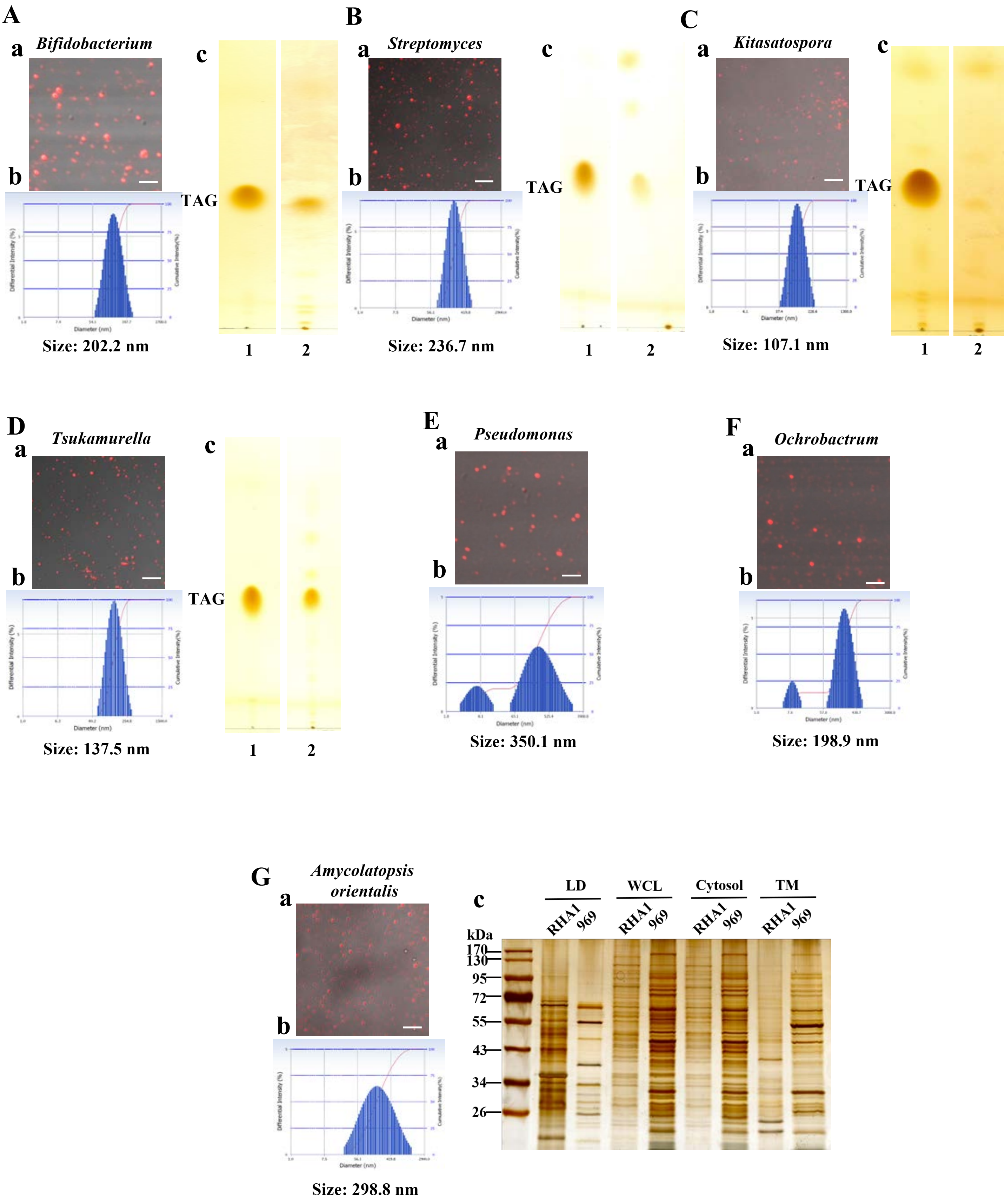

### Figure S4 Isolation and Analysis of Lipid Droplets from Bacteria

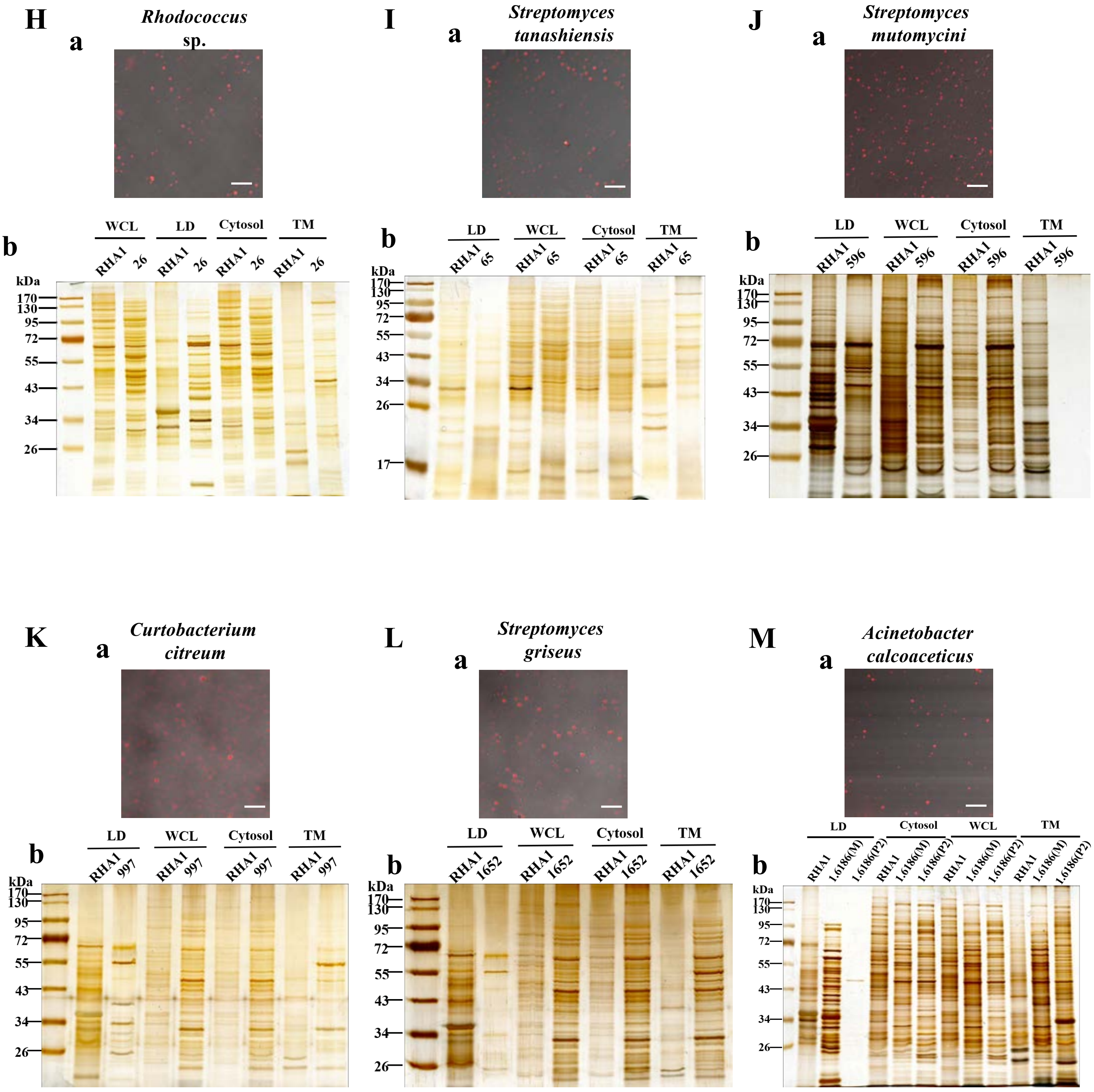

### Figure S4 Isolation and Analysis of Lipid Droplets from Bacteria

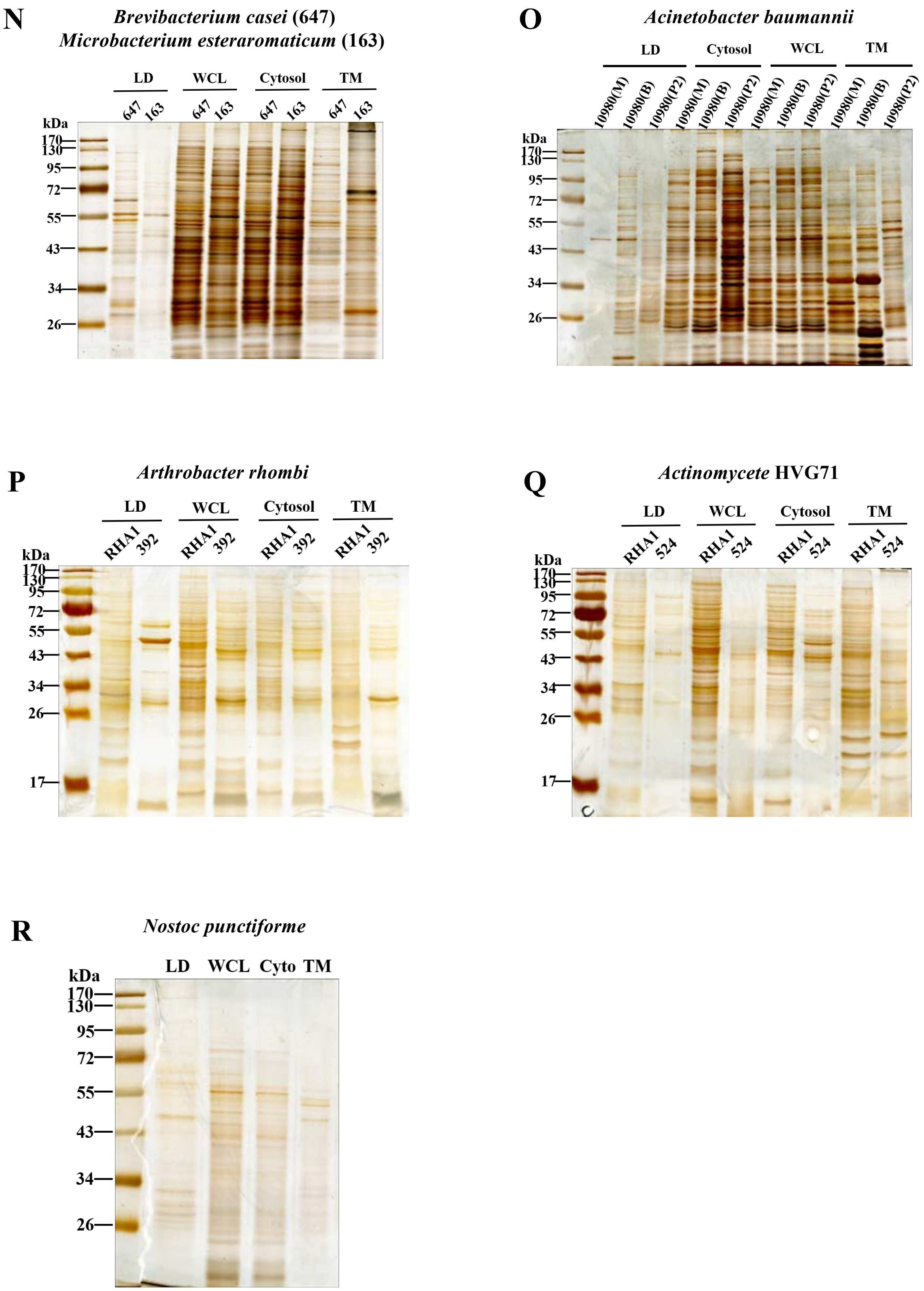

### Figure S5 Isolation and Proteomic Analysis of Lipid Droplets from Bacteria

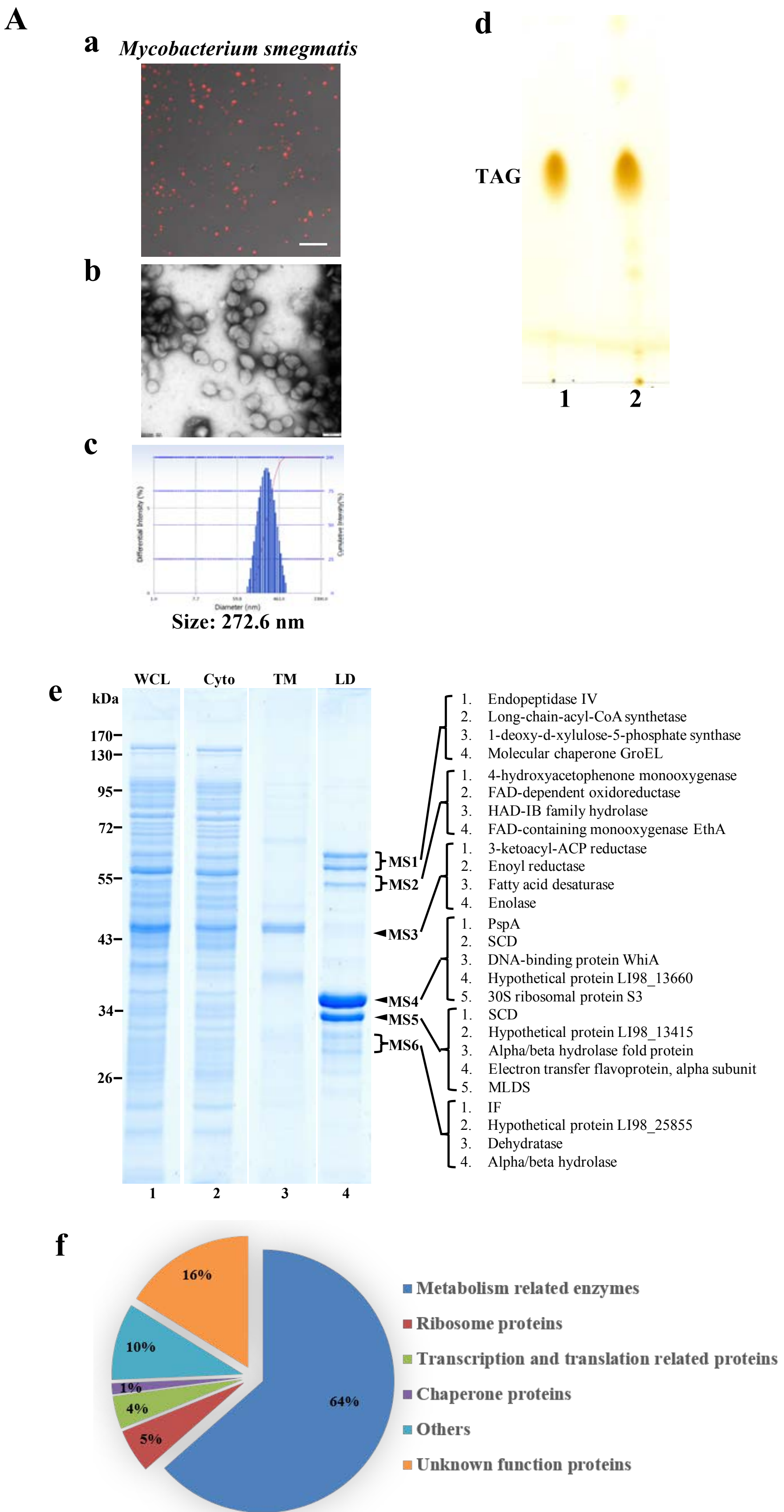

### Figure S5 Isolation and Proteomic Analysis of Lipid Droplets from Bacteria

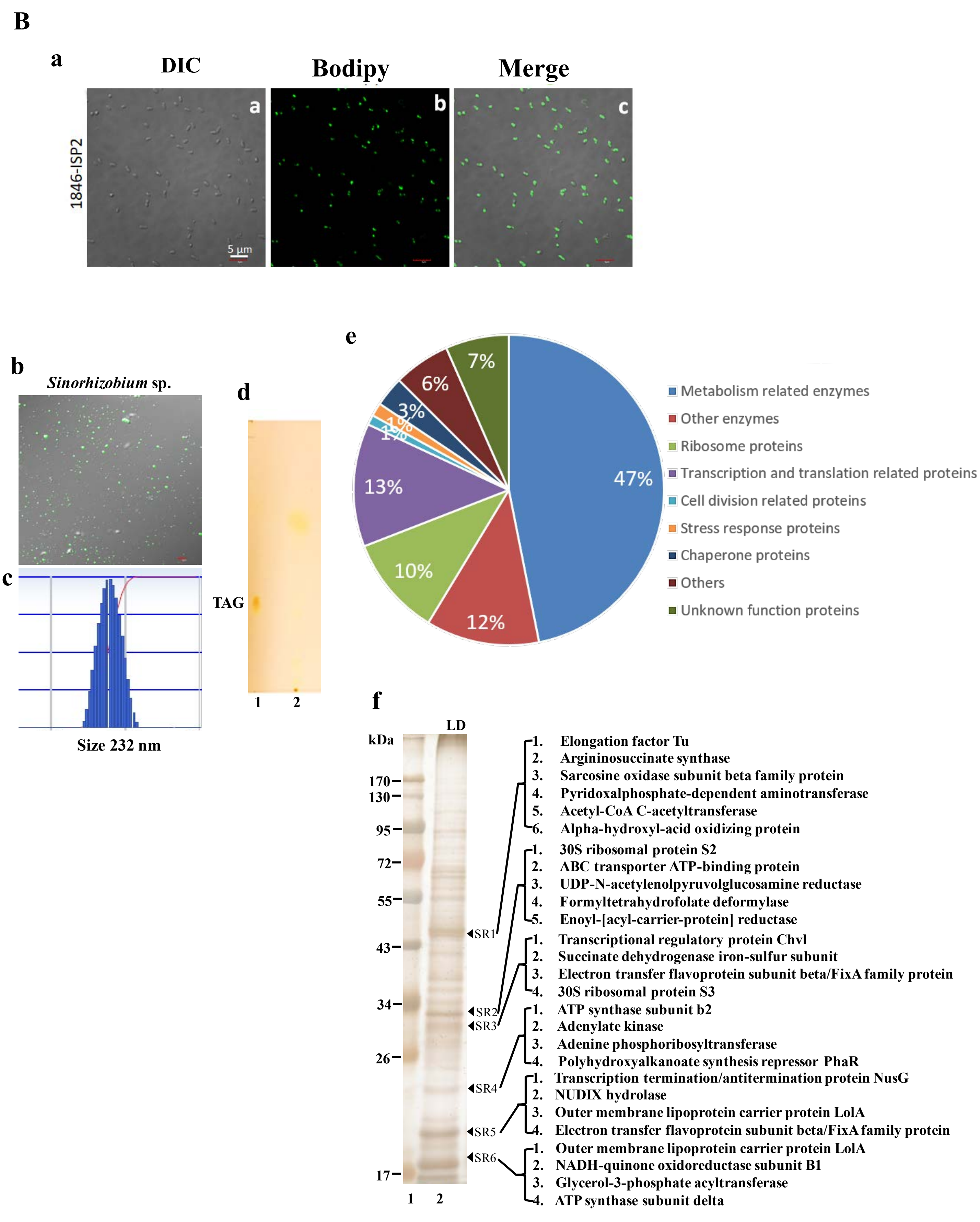

### Figure S5 Isolation and Analysis of Lipid Droplets from Bacteria

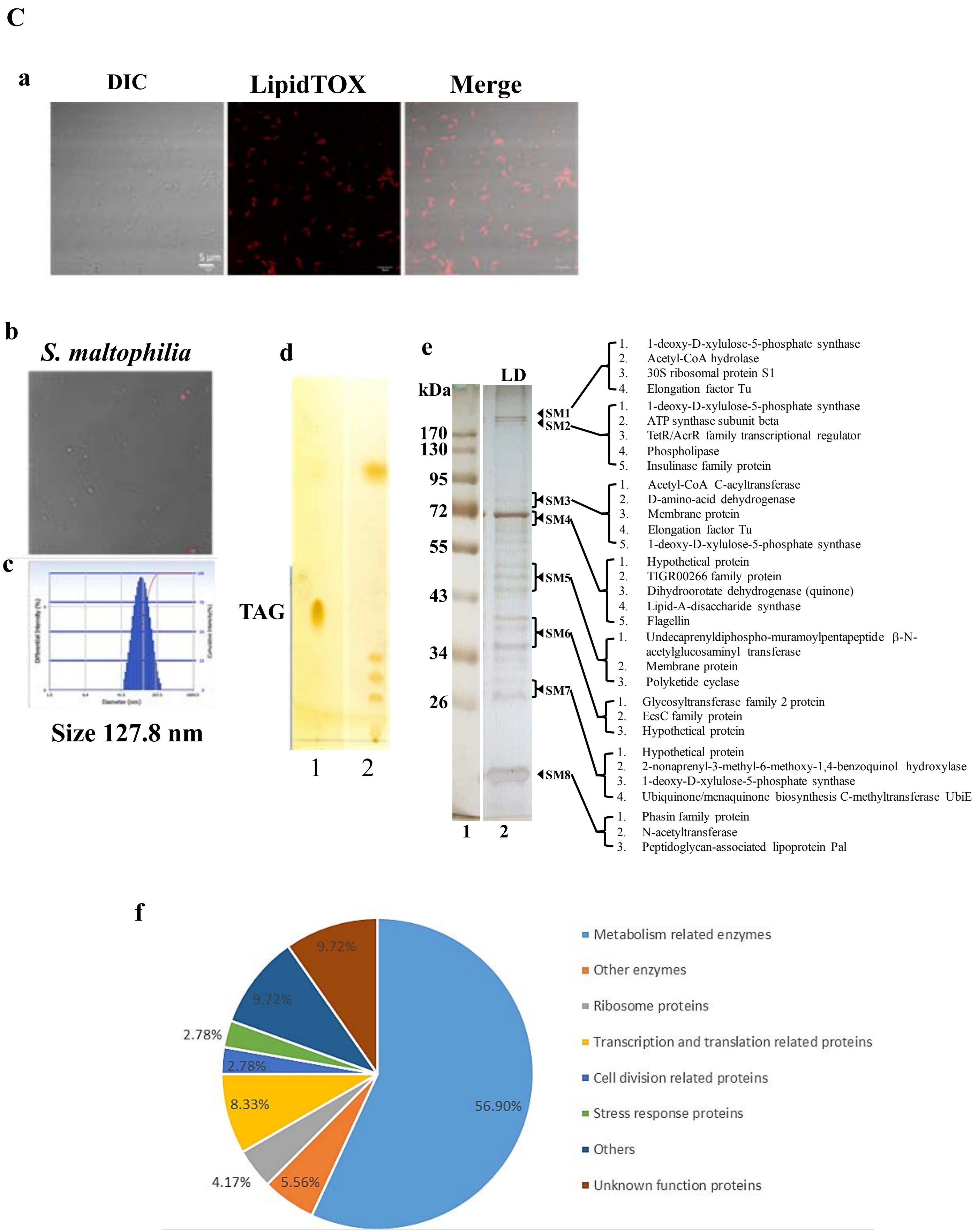

### Figure S6 Generation of Lipid Droplets by Engineered *E. coli*

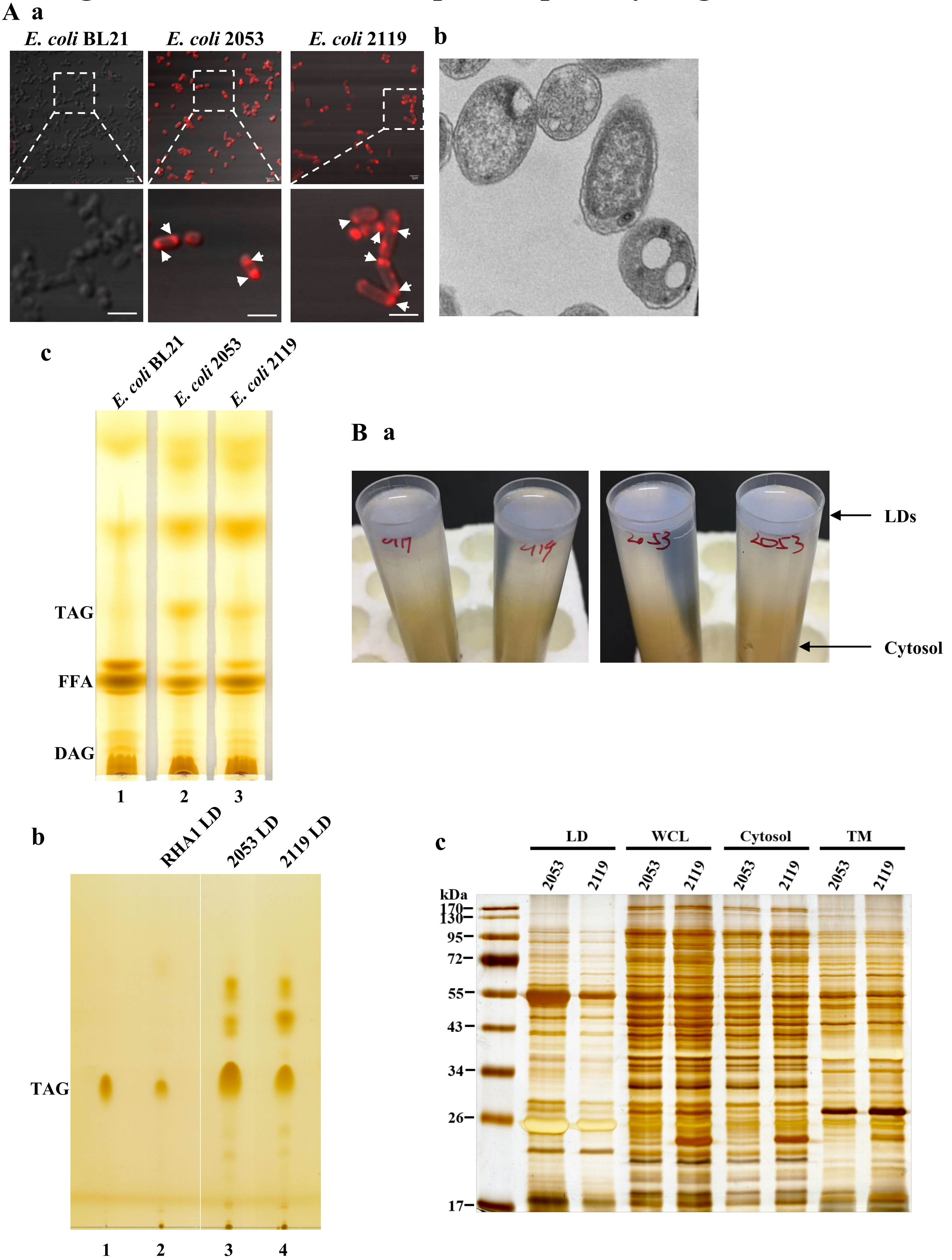

### Figure S7 PspA Is a Lipid Droplet Protein and Conserved in Bacteria

**A**

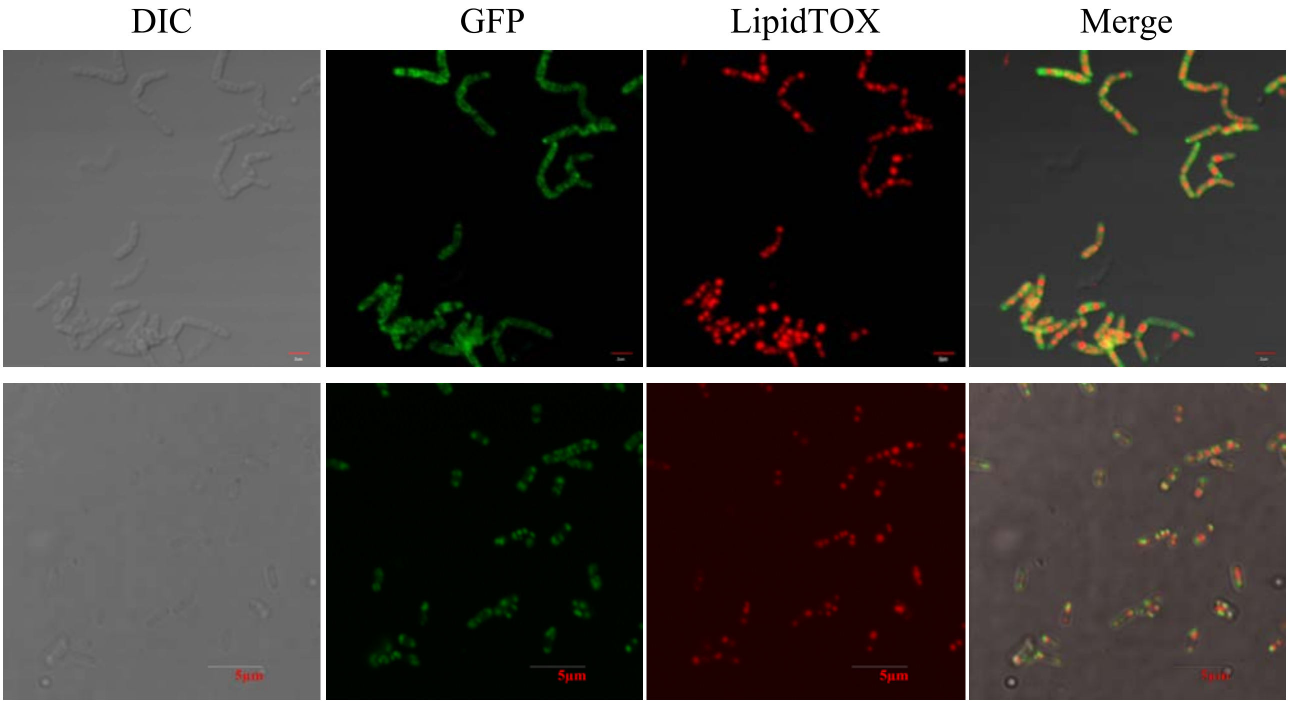


**B**

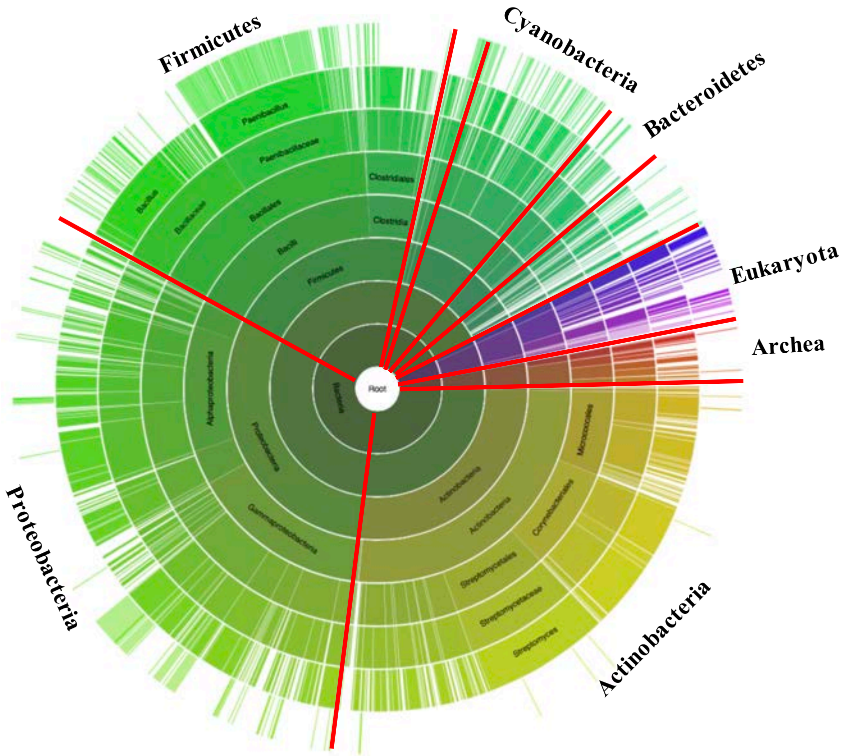

### Figure S8 Lipid Droplet Is an Ancient and Inheritable Organelle

A

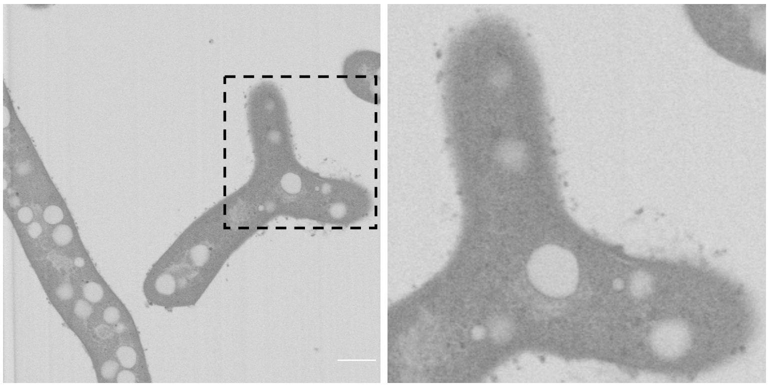

B

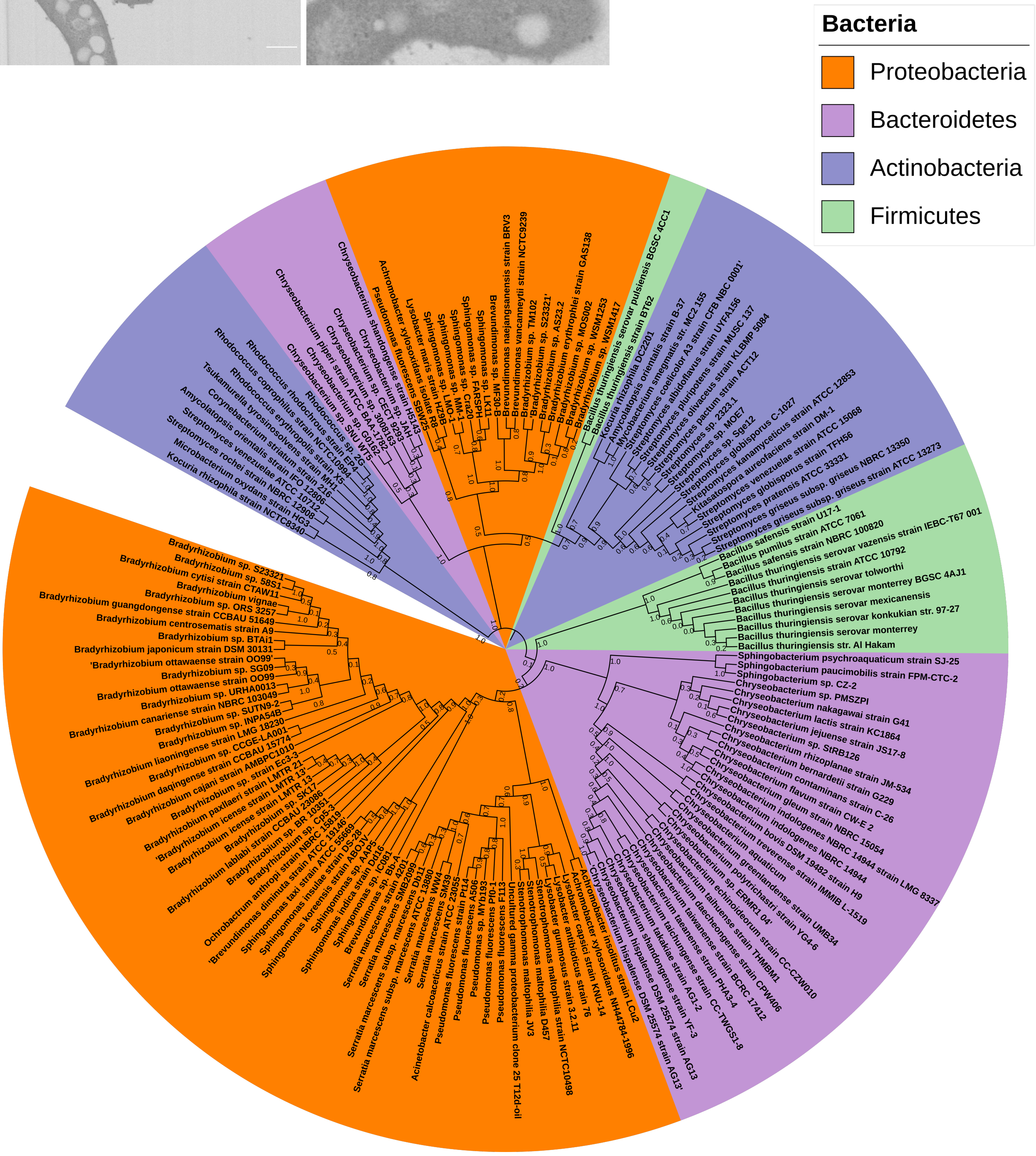
